# Virus Entry is a Major Determinant of HCMV Latency in Monocytes

**DOI:** 10.1101/2024.10.26.619803

**Authors:** Yaarit Kitsberg, Aharon Nachshon, Tamar Arazi, Karin Broennimann, Tal Fisher, Alexander Wainstein, Yaara Finkel, Noam Stern-Ginossar, Michal Schwartz

## Abstract

Human cytomegalovirus (HCMV) infection can result in either productive or latent infection, the latter enabling life-long viral persistence. Monocytes support latent infection but become permissive to productive infection upon differentiation into macrophages. The molecular basis for these differentiation-driven differences has been largely attributed to chromatin-mediated repression of the viral genome. Using metabolic labeling of newly synthesized RNA during the early stages of infection, we observed markedly lower viral transcription in monocytes compared to macrophages. Unbiased comparison of the two cell types revealed that this difference is partly due to reduced viral entry in monocytes: fewer viruses enter, and correspondingly fewer genomes reach the nucleus. Indeed, ectopic expression of known HCMV entry receptors in monocytes enhanced viral entry and consequently facilitated productive infection, demonstrating that these cells can support full replication if entry is efficient. We further identified integrin β3 as a surface protein upregulated upon differentiation that plays an important role in HCMV entry into macrophages, partially accounting for the observed differences in entry efficiency. Finally, we show that cells receiving fewer viral genomes are the ones that establish latent infection and have the capacity to reactivate. Overall, our findings reveal that entry is a previously unrecognized factor contributing to latent infection in monocytes, adding a critical layer to the paradigm of HCMV latency.

## Introduction

Human Cytomegalovirus (HCMV) is a human beta-herpesvirus infecting the majority of the population worldwide. Like other herpesviruses, HCMV persists through the lifetime of its host by establishing latency. Cells of the hematopoietic system were identified as key sites for HCMV latency. CD34+ hematopoietic stem cells (HSCs) and early progenitors of the myeloid system in the bone marrow as well as blood monocytes are the main cells in which HCMV latency has been characterized ^1–3^. In contrast, terminally differentiated myeloid cells such as macrophages and dendritic cells are considered permissive for productive HCMV infection, and differentiation of infected monocytes to these cell types can lead to viral reactivation ^4–7^.

We previously showed that when macrophages are infected, the main factor that dictates productive infection is the initial levels of viral gene expression and specifically the expression of HCMV immediate early genes, IE1 and IE2 ^8^. The accepted underlying assumption is that chromatin-dependent repression of the viral genome, and specifically of immediate early genes, is the basis for latent infection in monocytes and HSCs ^9^, and that differentiation leads to distinct chromatin deposition that enables viral gene expression ^10,11^. However, the molecular roots for these differences in repression upon differentiation are not fully understood.

Here, using metabolic labeling of newly synthesized mRNA, we reveal that compared to macrophages, early viral gene expression in undifferentiated monocytic cells is much lower. By systematically comparing monocytes and their differentiated counterparts, we show that this strong difference in gene expression is partially due to differences in viral entry and that in monocytes, less viruses enter the cells and correspondingly, less viral genomes reach the nucleus. Remarkably, ectopic expression of known HCMV entry receptors in monocytes results in more efficient viral entry and concomitantly, productive infection. We further show that integrin β3, which was previously associated with HCMV entry ^12^, is upregulated upon differentiation and plays a critical role in macrophage infection. Importantly, we demonstrate that infected cells that receive less viral genomes are the source of latent cells that can later reactivate. Overall, we uncover that inefficient entry, due to specific cell surface composition, is a major factor precluding productive infection in monocytes, and that this inefficient entry leads to cells receiving low levels of viral genomes resulting in the establishment of latent infection.

## Results

### The early phase of HCMV infection in monocytes is marked by strikingly low viral gene expression

Monocytes are known to be latently infected with HCMV and do not support productive infection, however, following differentiation they become permissive to productive infection ^4,5^. Indeed, we could recapitulate these differentiation-based differences in HCMV infection in primary monocytes isolated from peripheral blood as well as in the myeloid cell lines THP1 and Kasumi-3, which are commonly used as cell models to study HCMV latency (Fig. 1a). We infected these cell types with an HCMV TB40-E strain containing a GFP reporter (HCMV-GFP), which allows convenient quantification of productively infected cells ^13^. In the monocytic cells, GFP expression remained low (Fig. 1b, Fig. S1a) and correspondingly, no infectious progeny production was detected at 10 days post-infection (dpi) (Fig. 1c). Differentiation of infected monocytes at 7 dpi led to reactivation of the virus in a portion of the cells (Fig. S1b) indicating that they are indeed latently infected. In contrast, differentiation of primary, THP1 or Kasumi-3 monocytes to monocyte-derived macrophages prior to infection, resulted in a distinct population of cells that expressed high levels of GFP at 3 dpi, indicating productive infection (Fig. 1b, Fig. S1a) and indeed these cells produced infectious progeny (Fig. 1c).

**Fig. 1.**
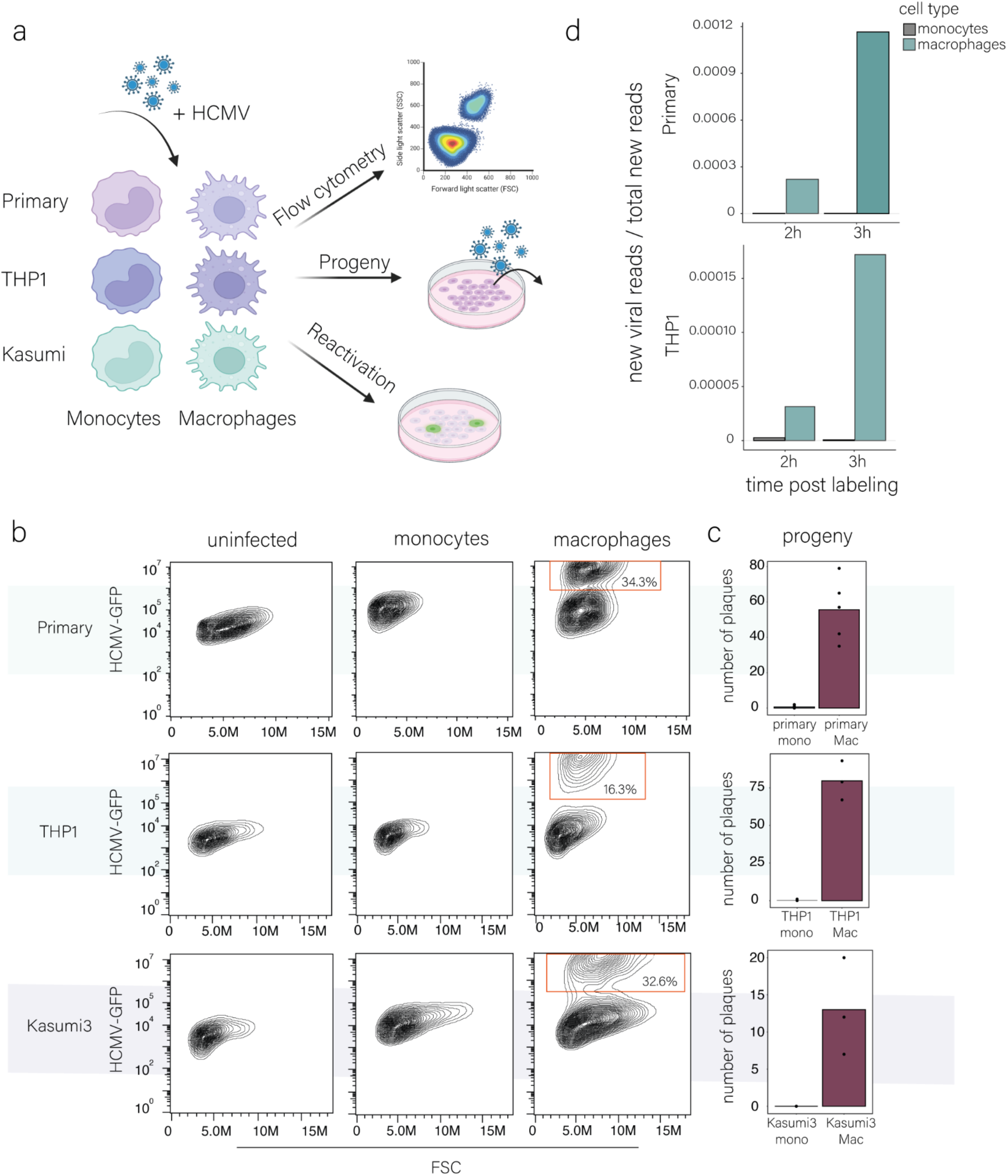
HCMV infection and viral gene expression in monocytes and macrophages. (a). Schematic illustration of the experimental setup. Monocytes and macrophages from primary, THP1 and Kasumi-3 cells were infected with HCMV-GFP. Infection levels, viral progeny, and reactivation were then analyzed. (b). Flow cytometry analysis of primary, THP1 and Kasumi-3 monocytes and their differentiated counterparts infected with HCMV–GFP (MOI =5). Analysis was performed at 3 days post-infection (dpi). The red gate marks the productive, GFP-bright cell population. FSC, forward scatter; M, million. (c). Measurements of infectious virus in supernatants collected from infected monocytes and macrophages at 10 dpi. Mono, monocytes; Mac, macrophages. n = 3-5. (d). Proportion of new viral reads out of the total new reads detected by SLAM-seq in infected primary and THP1 monocytes and macrophages. Infected cells were labeled with 4sU at 3 hpi and harvested for SLAM-seq after 2 hours (left bars) or 3 hours (adjacent right bars) of labeling.

We recently showed that initial levels of viral gene expression are a major factor in dictating productive infection in macrophages ^8^ and therefore wanted to accurately assess if differences in initial viral gene transcription can explain these substantial differences in infection. Since our work and that of others, demonstrated that during early monocyte infection newly transcribed RNA is masked by virion-associated input RNA ^14,15^, we aimed to directly measure early viral gene transcription. We applied thiol (SH)-linked alkylation for metabolic sequencing of RNA (SLAM-seq)^16^. SLAM-seq facilitates the measurement of newly transcribed RNA based on 4-thiouridine (4sU) incorporation into newly synthesized RNA. After RNA is extracted, 4sU is converted to a cytosine analog using iodoacetamide, and these U to C conversions are identified and quantified by RNA sequencing. We applied SLAM-seq to both primary and THP1 monocytes and macrophages, starting labeling at 3 hours post-infection (hpi) for two and three hours (cells were harvested at 5 and 6 hpi). We confirmed that 4sU labeling does not affect cell viability (Fig. S1c) and successfully generated quality SLAM-seq libraries in both primary and THP1-derived cells, with over 5,389 genes quantified and a high U-to-C conversion rate (Fig. S1d). The levels of newly synthesized cellular transcripts were comparable between infected primary monocytes and macrophages, as well as between infected THP1 monocytes and THP1 macrophages in both two and three hours of labeling, indicating there are no major biases in our labeling (Fig. S1e). Remarkably, newly synthesized viral transcripts were extremely low in monocytes regardless of the labeling time, while in macrophages new viral transcripts were detected after two hours of labeling and their relative fraction further increased at 3 hours of labeling (Fig. 1d). These newly synthesized viral transcripts in macrophages were predominantly immediate early genes (UL122 and UL123), reflecting the initiation of an infection cycle (Fig. S1f). These results demonstrate that already at very early stages of infection in monocytes (both primary and THP1), viral genes are weakly transcribed, while in macrophages they are efficiently expressed.

### Cell surface proteins are upregulated upon monocyte to macrophage differentiation

The immediate vast difference in the levels of synthesized viral transcripts in infected monocytes, compared to macrophages, indicates a major difference in HCMV’s ability to initiate gene expression in these two cell types. To unbiasedly search for candidate factors that may explain these dramatic differences, we performed RNA-seq on primary monocytes, THP1 monocytes and Kasumi-3 myeloid progenitor cells, as well as on macrophages derived from the same cells. Thousands of genes were differentially expressed upon differentiation (Table S1). Pathway enrichment analysis in each of the cell types (primary monocytes, THP1 and Kasumi-3 cells) revealed that common pathways change upon differentiation of THP1 and Kasumi-3 cells, while different pathways change upon differentiation of primary monocytes (Fig. 2a). In primary monocytes, the most significantly differential pathways were related to inflammation and innate immunity that mainly decreased upon differentiation, with interferon response pathways being the most significantly reduced (Fig. 2a). This is in line with our previous work showing that intrinsic expression of interferon stimulated genes (ISGs) is decreased upon differentiation, and that this reduction contributes to the increased susceptibility of macrophages compared to monocytes ^8^. In THP1 and Kasumi-3, which are tumor-derived cell lines, the most significant changes were a decrease in pathways related to cell proliferation, such as E2F targets, Myc targets and G2/M checkpoint (Fig. 2a), in line with previous reports of a significant reduction in these cells’ proliferation capacity upon differentiation ^17,18^. The effect of cell proliferation on HCMV infection has been extensively characterized in fibroblast infections ^19,20^. These findings suggest that in THP1 and Kasumi-3, differences in permissivity following differentiation may be partially attributed to reduction in proliferative capacity. Significantly enriched pathways that were shared between all three cell types included a reduction in apical junction and in myogenesis, both pathways not intuitively related to myeloid differentiation processes or to early viral gene expression. Parsimoniously, we expect the same mechanism to explain the difference in infection upon differentiation in all three cell types. Thus to reveal molecular processes relevant for the different infection outcomes, we focused on common differentially expressed genes across cell types. Upon differentiation, 213 genes were commonly upregulated between primary, THP1 and Kasumi-3 cells, which is a significant overlap (*p* < 10^-5^, Fig. S2a), and 42 genes were commonly downregulated in all three cell types (*p* = 0.133, Fig. S2a). Pathway enrichment analysis on the commonly upregulated genes yielded several pathways related to differentiation and maturation of immune cells and immune signaling (Fig. S2b). Since we revealed massive differences in initial viral gene expression, we focused on processes that can potentially explain these differences. Although there is a major focus in the field on chromatin-related factors that regulate HCMV repression in monocytes ^21,22^, such factors, as a group, were not significantly enriched in the shared genes (Fig. S2c). Nevertheless, four chromatin-related factors were downregulated upon differentiation in the three cell types, including CHD3, which is implicated in the repression of the HCMV genome through the recruitment of HDACs^23,24^. We therefore explored the potential involvement of histone deacetylation, which is reported to play a key role in the repression of viral genes during HCMV latency ^25,26^. We tested the ability of the potent HDAC inhibitor, TrichostatinA (TSA), which is known to induce expression of IE (immediate early) genes in THP1 cells ^27^, to induce productive infection in monocytes. While TSA treatment indeed led to an increase in the percentage of productively infected monocytes, a comparable effect was also observed in macrophages. Furthermore, the TSA effect was small compared to the effect of differentiation, suggesting that additional factors likely contribute to the differences between monocytes and macrophages (Fig. 2b and Fig. S2d).

**Fig. 2.**
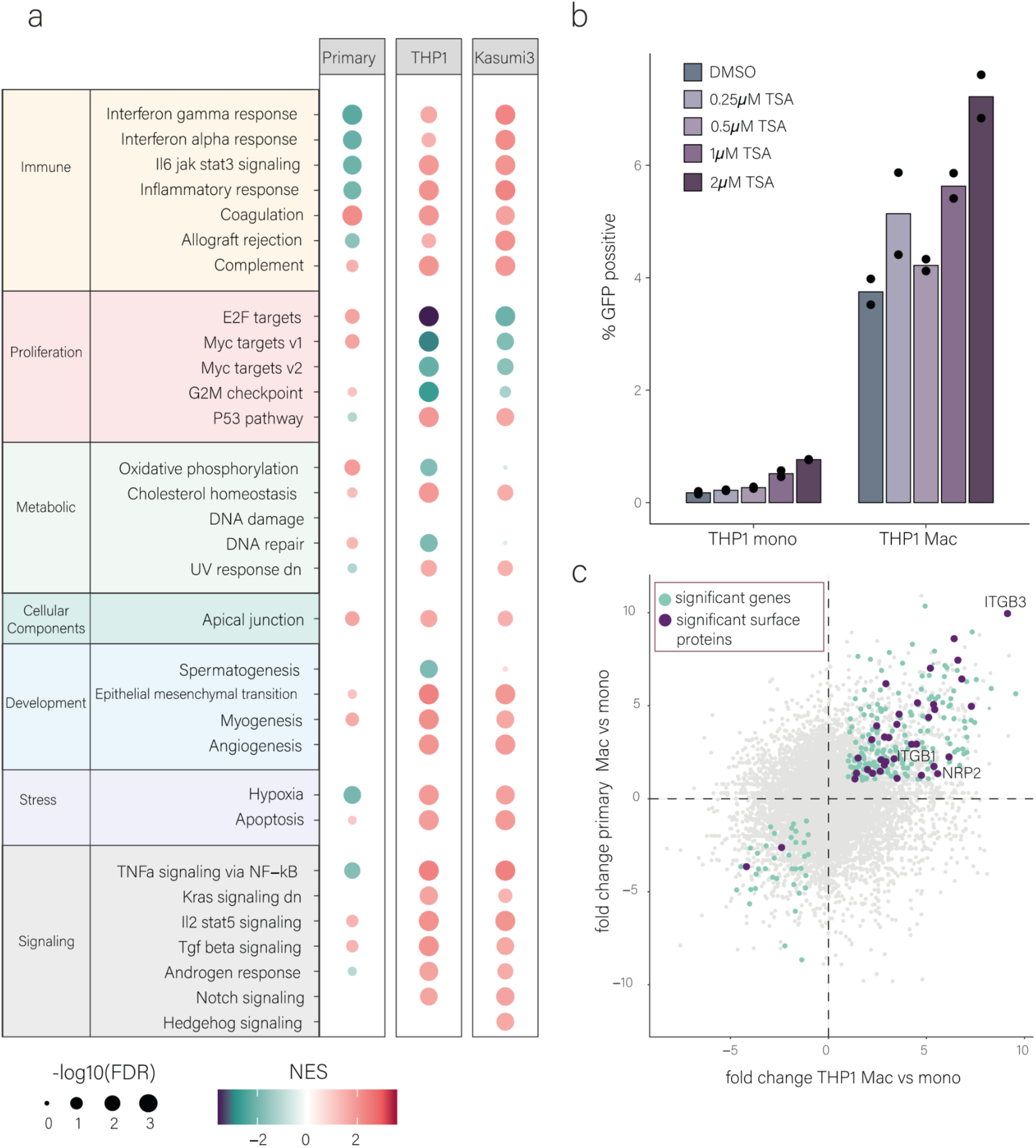
Changes in gene expression upon differentiation. (a). Summary of Hallmark pathway enrichment analysis of differentially expressed genes upon monocyte to macrophage differentiation in primary, THP1 and Kasumi-3 cells. FDR, false discovery rate; NES, normalized enrichment score. (b) THP1 monocytes and macrophages were treated with increasing concentrations of TSA. Cells were treated with TSA or DMSO control at 5 hpi and analyzed at 3 dpi by flow cytometry. n=2. Gating strategy is shown in Fig. S1a. (c) Scatterplot of the fold change (FC) from RNA-seq data between primary monocytes and macrophages, relative to the fold change of THP1 monocytes and macrophages. Light blue dots mark significantly changing genes (FDR < 0.05, LFC > 1) in all three cell types. Dark purple dots mark single transmembrane genes that are significantly changing in all three cell types (P = 0.042). Names of significantly changing cell surface proteins in all three cell types involved in HCMV entry compiled based on ^29^ and other studies (see Table S2) are shown.

Intriguingly, we observed a significant upregulation of cell surface proteins as a group, following monocyte-to-macrophage differentiation (p = 0.042; Fig. 2c), suggesting substantial remodeling of the cell surface composition. Given that such changes can influence viral entry and, consequently, viral gene expression, we asked whether surface proteins previously implicated in HCMV entry are among the upregulated genes. Indeed, HCMV-associated entry factors were significantly enriched among the commonly upregulated surface proteins (p = 0.0062, hypergeometric enrichment test; Table S2, Fig. 2c). These upregulated genes include NRP2, which mediates HCMV entry into non-fibroblasts cells (through the viral pentamer entry complex)^28^ as well as two integrins, ITGB1 and ITGB3, which were shown to play a role in HCMV entry ^29^ (Fig. 2c and Table S2). These changes in the expression of cell surface proteins involved in HCMV entry, pointed to possible unexplored differences in viral entry between monocytes and macrophages.

### Inefficient viral entry into monocytes is a major cause for low viral gene expression

To explore if indeed disparities in viral entry may explain some of the differences in viral gene expression, we quantified the number of viral genomes in the nuclei of infected primary and THP1 monocytes and macrophages by DNA FISH. At 12 hpi, we could detect viral genomes in the nuclei of monocytes, but their amount was much lower than the amount of viral genomes in the nuclei of macrophages (Fig. 3a, S3, 3b, Supplementary Video 1). To further examine differences in entry efficiency, we utilized a virus in which the tegument protein UL32 is tagged with GFP (UL32-GFP^30^), allowing fluorescent tracking of viral particles. In correspondence with our DNA-FISH measurements, we found significantly more viral particles within infected macrophages compared to infected monocytes (Fig. 3c and 3d). These results suggest there is a considerable difference in the efficiency of viral entry between monocytes and macrophages, with less viral genomes reaching the nucleus of monocytes.

**Fig. 3.**
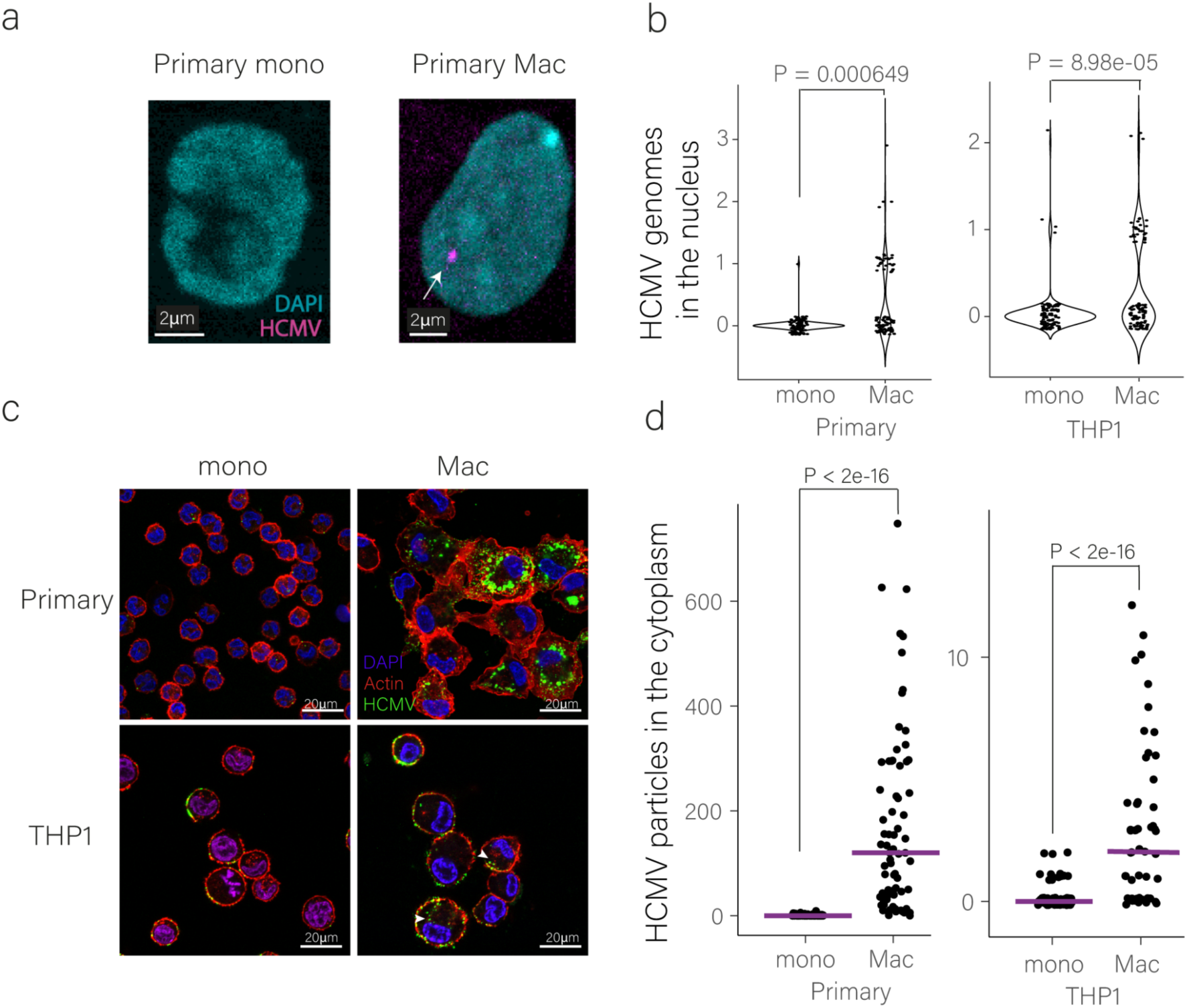
HCMV genomes are detected at very low levels in the nucleus and cytoplasm of infected monocytes compared to macrophages. (a). Images of infected primary monocyte and macrophage nuclei at 12 hpi. (MOI=5). The HCMV genome was probed using DNA-FISH. (b). Quantification of viral genomes detected in the nuclei of infected primary or THP1 monocytes (n=87 and n=112, respectively) and macrophages (n=93 and n=109, respectively) by DNA-FISH at 12 hpi. The P-value was calculated using Poisson regression. (c). Images of HCMV particles (labeled by UL32-GFP) in infected primary and THP1 monocytes and macrophages at 1 hpi (MOI=5). Actin staining was used to visualize cell borders and DAPI for the nuclei. (d). Quantification of viral particles within the cytoplasm of infected primary and THP1 monocytes (n=115 and n=78, respectively) and macrophages (n=71 and n=53, respectively) at 1 hpi (presented in c). Viral particles were counted using FIJI image processing and statistical analysis was performed using Poisson regression. Mono, monocytes; mac, macrophages.

To test whether inefficient viral entry contributes to non-productive infection of monocytes, we aimed to increase viral entry efficiency and test the effect on infection. To this end we ectopically expressed PDGFRα, a well characterized entry receptor of HCMV in fibroblasts^31^, which is not expressed in either monocytes or macrophages (Fig. S4a), in THP1 monocytes (THP1-PDGFRα, Fig. S4b and c). Remarkably, infection of THP1-PDGFRα with HCMV-GFP, resulted in a distinct population of GFP-bright cells at 3 dpi, indicating these cells are productively infected (Fig. 4a). Furthermore, productively infected cells were those with higher surface expression levels of PDGFRα suggesting a direct connection between efficient entry and the ability to establish productive infection (Fig. 4b). Infected THP1-PDGFRα supported viral genome replication (Fig. 4c) and generated viral replication compartments (Fig. S4d). However, viral titers were extremely low (Fig. S4e), and as shown below this is likely due to the ectopically expressed PDGFRα interfering with the infectivity of viral progeny.

**Fig. 4.**
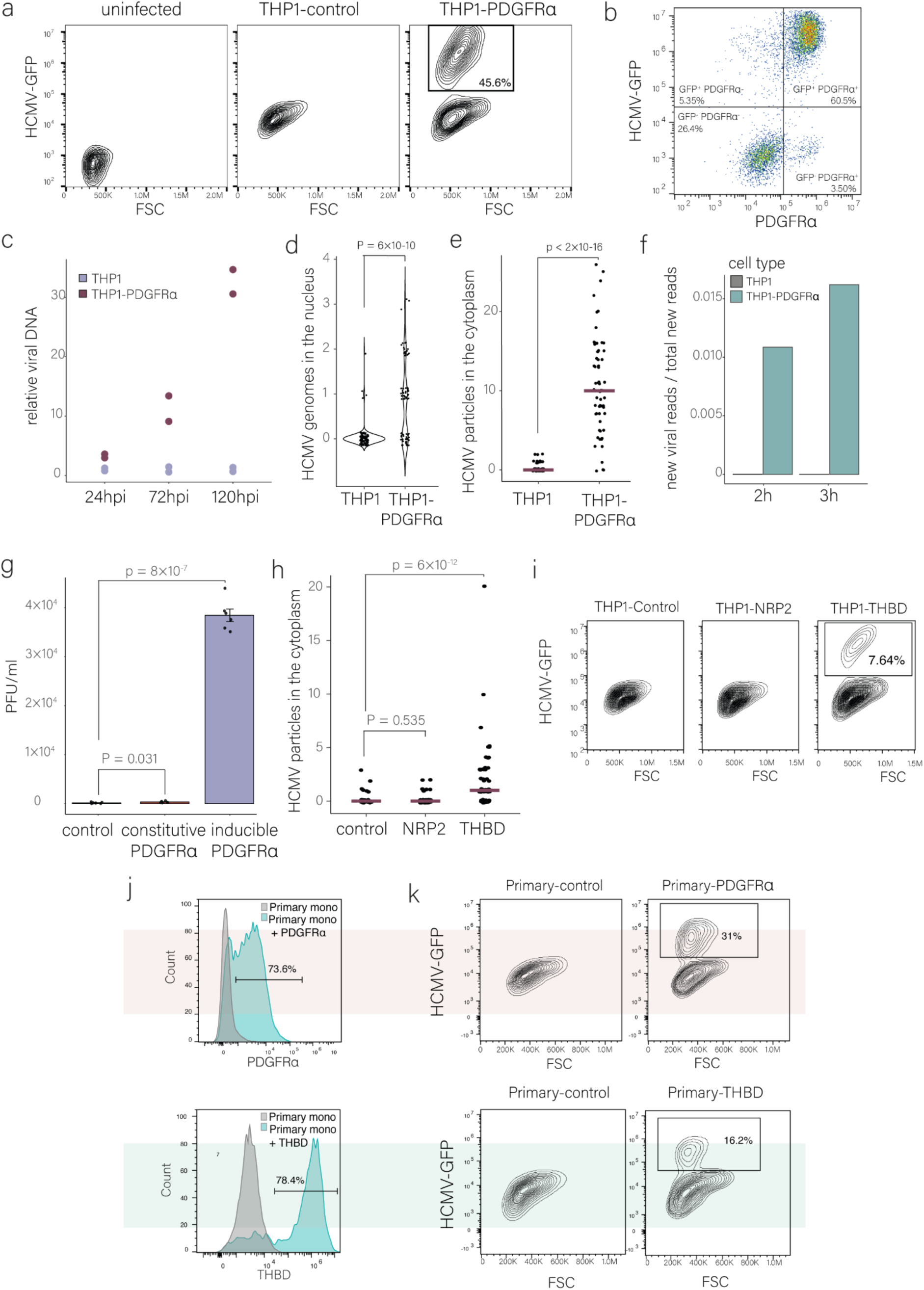
Ectopic expression of known HCMV entry receptors in monocytes leads to productive infection. (a). Flow cytometry analysis of THP1 and THP1 overexpressing PDGFRα (THP1-PDGFRα) infected with HCMV–GFP (MOI=5) at 3 dpi. The gate marks the productive, GFP-bright cell population. Gating strategy is shown in Fig. S1a. (b). Flow cytometry analysis of THP1 overexpressing PDGFRα (THP1-PDGFRα) infected with HCMV–GFP (MOI=5) at 3 dpi showing HCMV-GFP level versus PDGFRα surface level. Gating strategy is shown in Fig. S1a. (c). Relative viral DNA levels in infected THP1 and THP1-PDGFRα at 24, 72 and 120 hpi. Viral DNA levels were measured by real-time PCR and were normalized to a cellular genomic target. n=2. (d). Quantification of the number of viral genomes in the nuclei of infected THP1 and THP1-PDGFRα monocytes at 12 hpi as detected by DNA-FISH (n=112 and n=84, respectively). P-value was calculated using Poisson regression. (e). Quantification of the number of viral particles within the cytoplasm of infected THP1 and THP1-PDGFRα monocytes using HCMV-UL32-GFP (MOI=5) at 1.h.pi. (n=78 and n=62, respectively). Viral particles were counted using FIJI image processing and statistics was performed using Poisson regression. (f). Proportion of new viral reads out of the total new reads detected by SLAM-seq in infected THP1 and THP1-PDGFRα monocytes. Infected cells were labeled with 4sU at 3 hpi and harvested for SLAM-seq after two (left bars) or three (adjacent right bars) hours of labeling. (g). THP1 monocytes constitutively expressing PDGFRα or induced to expressed PDGFRα 24h before infection were infected with HCMV-GFP compared to control infected cells. At 8 dpi, viral supernatant was used to infect recipient wild-type fibroblasts. Forty-eight hours later, the percentage of GFP-positive recipient cells was determined by flow cytometry and used to calculate the number of plaque-forming units (PFU). p-value was calculated using a two-sided student t-test, n = 6. (h). Quantification of the number of viral particles, using HCMV-UL32-GFP (MOI=5), within the cytoplasm of infected THP1 monocytes with induced expression of mCherry as a control, NRP2 or THBD at 1.h.pi. (n=50, 67 and 63, respectively). (i). Flow cytometry analysis of THP1 monocytes overexpressing control, THBD and NRP2, using a dox-inducible system, infected with HCMV– GFP (MOI=5) at 3 dpi. The gate marks the productive, GFP-bright cell population. Gating strategy is shown in Fig. S1a. (j). Flow cytometry analysis of PDGFRα surface expression on primary monocytes transfected with control, THBD and PDGFRα mRNA at 12h after transfection. Gating strategy is shown in Fig. S1a. (k). Flow cytometry analysis of primary monocytes transfected with PDGFRα, THBD or control mRNA for 12h before HCMV infection (MOI=5). Cells were analyzed at 3dpi. Gating strategy is shown in Fig. S1a.

To substantiate that ectopic expression of PDGFRα in THP1 monocytes directly affects viral entry, we quantified viral genomes at 12 hpi and found that in contrast to the parental THP1, in THP1-PDGFRα viral genomes reach the nucleus in a considerable portion of the cells (Fig. 4d, Supplementary Video 1). Furthermore, infected THP1-PDGFRα monocytes had significantly more viral particles in the cytoplasm than THP1 monocytes (Fig. 4e), resembling the levels observed in THP1-derived macrophages (Fig. S4f). Correspondingly, these cells transcribe viral genes at 5 hpi, as measured by SLAM-seq (Fig. 4f). Notably, differentiation of THP1-PDGFRα cells further enhanced viral entry, suggesting that, as expected, differentiation facilitates HCMV entry through a PDGFRα-independent mechanism (Fig. S4f). Importantly, overexpression of PDGFRα did not result in differentiation of the cells, as the cells did not differ from the parental THP1 cells morphologically or in their expression levels of macrophage-related surface markers (Fig. S4g and Fig. S4h). We further performed transcriptomic as well as proteomic analyses on THP1-PDGFRα and the parental cells and found only minor changes in gene expression and protein composition, all of which are not related to macrophage differentiation or innate immunity (Fig. S4i, S4j and Tables S3), negating the possibility of indirect effects of PDGFRα expression. To further rule out indirect effects of long-term PDGFRα expression, we repeated the experiment with an inducible expression system in which PDGFRα expression is induced shortly prior to infection (Fig. S4k). Also in these conditions, PDGFRα expression led to productive infection in a substantial percentage of cells (Fig. S4l). Notably, when using the inducible PDGFRα system to only transiently express PDGFRα (Fig. S4k), infected THP1 monocytes produced infectious viral progeny. This confirms that a transient increase in HCMV entry in THP1 monocytes supports completion of the viral replication cycle, and suggests that constitutive PDGFRα expression likely interferes with particle infectivity (Fig. 4g). Indeed, when we examined the amount of viral genomes in the supernatant by qPCR we found that when PDGFRα is expressed, either by a constitutive or an inducible promoter, there are much more viral genomes compared to control cells (Fig. S4m). These findings support the hypothesis that constitutive PDGFRα expression in THP1-PDGFRα cells compromises the infectivity of the virions they release. Finally, we treated THP1-PDGFRα cells with imatinib which blocks PDGFRα signaling ^32^ and found that imatinib did not affect productive infection suggesting that PDGFRα effects on infection are independent of its signaling and are likely mostly mediated by changes in viral entry (Fig. S4n) in line with previous reports ^31,33^.

We next explored whether additional HCMV entry receptors can induce productive infection. Using the inducible system in THP1 monocytes, we over-expressed two additional established receptors of HCMV, NRP2 and THBD ^28,34,35^ (Fig. S4o), and then infected these cells with HCMV-GFP. While overexpression of NRP2 did not affect HCMV entry, overexpression of THBD led to a significant increase in viral entry into monocytes (Fig. 4h). Correspondingly, overexpression of THBD led to a distinct population of productively infected monocytes, as evident from high viral GFP expression as well as production of infectious viral progeny (Fig. 4i and Fig S4p), while NRP2 overexpression did not affect infection (Fig. 4i). Productively infected cells were those with higher surface expression levels of THBD suggesting a direct connection between efficient entry and the ability to establish productive infection (Fig. S4q). THBD overexpression in THP1 monocytes did not induce differentiation, as they did not differ from the parental THP1 cells morphologically (Fig. S4g) or in their expression levels of macrophage-related surface markers (Fig. S4h). Transcriptomic analysis revealed only minor changes in gene expression, all of which are not related to macrophage differentiation or innate immunity (Fig. S4r and Table s4), negating the possibility of indirect effects of THBD expression.

To examine if this effect of enhancing viral entry can also be recapitulated in primary monocytes, we transfected primary monocytes with in-vitro transcribed PDGFRα mRNA (Fig. 4j). Infection of these cells with HCMV-GFP resulted in a distinct population of GFP-bright cells at 3 dpi (Fig. 4k). Although extensive cell death (likely resulting from transfection-related stress) prevented analysis of viral progeny, we detected robust viral genome replication (Fig. S4s). Similarly, transfection of primary monocytes with THBD RNA (Fig. 4j) resulted in a distinct population of GFP-bright cells (Fig. 4k). These results indicate that increasing viral entry in primary monocytes facilitates initiation of viral gene expression and viral genome replication indicating productive infection is likely taking place.

### Integrin *β*3 plays a role in HCMV entry into macrophages

Our findings indicate that differences in viral entry efficiency contribute to the different outcomes of HCMV infection in monocytes and macrophages but the source to these differences in entry remains unclear. We analyzed the potential involvement of several cell surface receptors implicated in HCMV infection which are upregulated upon differentiation. First we focused on NRP2, NRP2 transcript levels increase upon differentiation of monocytes to macrophages in the three cell types tested (Fig. 5a). Indeed, cell surface staining illustrated NRP2 is not expressed on the surface of monocytes and differentiation to macrophages was accompanied by low but detectable cell surface expression (Fig. 5b). Although we found that NRP2 ectopic expression in monocytes is not sufficient for inducing productive infection in monocytes (Fig. 4i), this could be due to the absence of additional factors which act together with NRP2 to facilitate entry. To test whether NRP2 is necessary for viral entry into macrophages, we used CRISPR-Cas9 to generate THP1 cells in which NRP2 is knocked out, which resulted in a partial knockout (Fig. S5a). We therefore also used siRNA to knockdown NRP2 expression (Fig. S5b). Infection of differentiated cells in which NRP2 was knocked out or down showed similar levels of infection compared to control cells (Fig. 5c-d and S5c-d), indicating NRP2 likely does not mediate HCMV entry into macrophages. To verify that our results are not impacted by possible mutations acquired during viral propagation, we sequenced the virus we used for infection (TB40 strain), and ruled out accumulation of mutations in the genes encoding the viral entry receptors (see methods section).

**Fig. 5.**
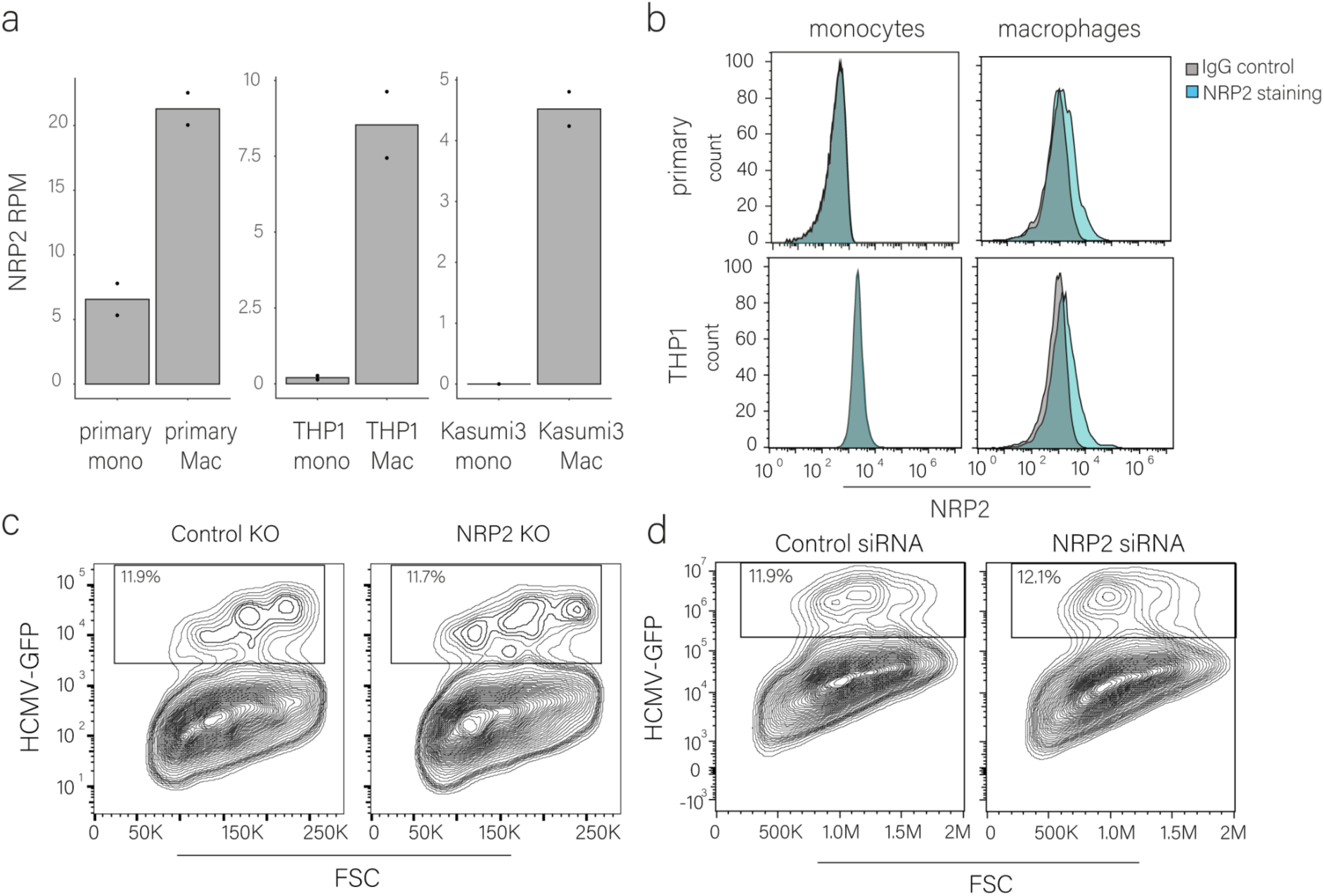
NRP2 does not facilitate HCMV entry into macrophages. (a). NRP2 expression in primary, THP-1 and Kasumi-3 monocytes and macrophages as measured by RNA-seq. Mono, monocytes; mac, macrophages; RPM, reads per million. n=2. (b). Flow cytometry analysis of NRP2 versus IgG control cell surface staining in primary and THP1 monocytes and macrophages. Gating strategy is shown in Fig. S1a. (c). Flow cytometry analysis of THP1 macrophages with NRP2 or control CRISPR knockout, infected with HCMV-GFP (MOI=5). Analysis was performed at 3 dpi. Gating strategy is shown in Fig. S1a. (d). Flow cytometry analysis of THP1 macrophages, transfected with NRP2 and control siRNA two days before infection with HCMV-GFP (MOI=5). Analysis was performed at 3 dpi. Gating strategy is shown in Fig. S1a.

We also analyzed two integrins, ITGB3, which encodes integrin β3 and ITGB1, which encodes integrin β1, both were significantly transcriptionally upregulated upon differentiation (Fig. 6a). These integrin β subunits can dimerize with different α subunits to form canonical heterodimers, some of which were implicated in HCMV entry ^12,36,37^. In agreement with our RNA-seq measurements, β3 surface expression was not detected in either primary or THP1 monocytes whereas in macrophages its expression was pronounced (Fig. 6b); β1 was expressed on the surface of monocytes but its expression significantly increased in macrophages (Fig. 6c and Fig. S6a, P < 0.0001 two-way ANOVA with Interaction).

**Fig. 6.**
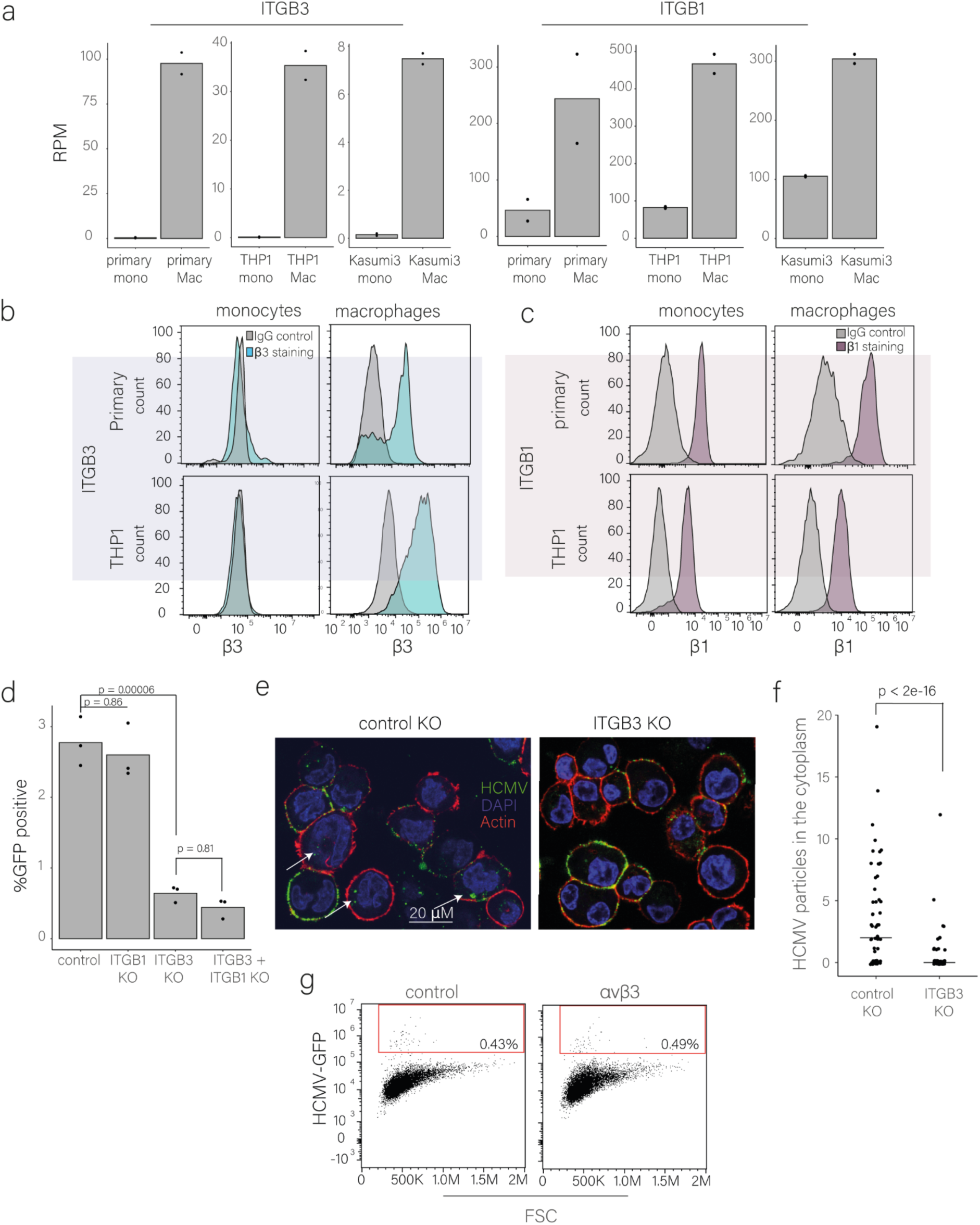
HCMV entry into macrophages is mediated through ITGB3. (a). ITGB3 and ITGB1 expression in primary, THP1 and Kasumi-3 monocytes and macrophages as measured by RNA-seq. RPM, reads per million. n=2. (b-c). Flow cytometry analysis of integrin β3 (b) and integrin β1 (c) cell surface levels in primary and THP1 monocytes and macrophages. Gating strategy is shown in Fig. S1a. (d). Flow cytometry analysis of control, ITGB3, ITGB1 and ITGB3 + ITGB1 knockout (KO) in THP1 macrophages infected with HCMV–GFP (MOI=5). Cells were analyzed at 3 dpi, p-value was calculated using a two-sided student t-test. n=3. Gating strategy is shown in Fig. S1a. (e-f). Representative Microscopy images (e) and quantification of viral particles using FIJI analysis software (f) of THP1 macrophages with ITGB3 knockout (KO) versus control knockout (n=67 and n=60, respectively) infected with HCMV-UL32-GFP (MOI=5) and imaged at 1.h.pi. Actin staining was used to visualize the cells’ borders and DAPI for nuclei staining. statistics was performed using Poisson regression. (g). Flow cytometry analysis of infected THP1 monocytes overexpressing ITGB3 and ITGAV (αvβ3), compared to mCherry control. Overexpression was induced for 24 hours using doxycycline prior to HCMV-GFP (MOI=5) infection. Cells were analyzed at 3 dpi. Gating strategy is shown in Fig. S1a.

To dissect if these integrins play a role in HCMV entry into macrophages, we generated CRISPR knockouts of either ITGB3 or ITGB1 in THP1 cells (Fig. S6b). Knockout of ITGB1 did not affect HCMV productive infection in THP1-derived macrophages. However, in the absence of ITGB3, differentiated macrophages were much less susceptible to productive infection compared to control cells (Fig. 6d and S6c). We also tested the knockout effect of both ITGB3 and ITGB1 but observed no cumulative effect beyond the effect of ITGB3 knockout (Fig. 6d and S6c).

We further validated these results by performing siRNA knockdown of ITGB3 in THP1 macrophages (Fig. S6d), showing that also knockdown leads to a significant decrease in productive infection of macrophages (Fig. s6e and s6f). By infecting with a UL32-GFP virus, we further show that in ITGB3 knockout macrophages viral entry was significantly reduced (Fig. 6e and 6f). These results illustrate that the expression of ITGB3 significantly increases upon monocyte to macrophage differentiation and that this increase plays a role in facilitating HCMV entry and subsequently in promoting infection of macrophages.

We next tested whether ectopic expression of ITGB3 alone is sufficient to promote productive infection in monocytes. However, ITGB3 overexpression did not enhance infection compared to control cells (Fig. S6g-h). Given that ITGB3 functions in complex with ITGAV (integrin αV), which is also upregulated during monocyte differentiation (Table S2), we co-expressed ITGB3 and ITGAV in THP1 monocytes using an inducible promoter (Fig. S6i, j). After sorting for double-positive cells (Fig. S6j), we infected them with HCMV-GFP. Co-expression of ITGB3 and ITGAV (αVβ3) likewise failed to increase productive infection relative to control cells (Fig. 6g, S6k). To verify the functionality of ectopically expressed ITGB3, we complemented ITGB3 KO and control macrophages by overexpressing ITGB3 (resistant to CRISPR editing) and we show that indeed complementation occurs and infection is restored, and moreover, overexpression of ITGB3 in macrophages further increases productive infection (Fig. S6l). These results demonstrate that, although ITGB3 plays a role in HCMV entry into macrophages, its increased expression in macrophages compared to monocytes is not the sole factor that facilitates the changes in viral entry and other proteins that are induced upon differentiation are likely required.

### Latency is established in cells receiving lower viral load

Our results suggest a hypothesis by which the number of incoming viral genomes plays a major role in determining the likelihood of productive infection. When the number of incoming viral genomes is low, productive infection is unlikely and instead latent infection may ensue. To test this, we aimed to isolate infected cells with lower load of viral genomes, which we hypothesized corresponds to infected cells that fail to initiate immediate early viral gene expression. We then sought to determine whether these cells are indeed latently infected (Fig. 7a). To directly assess this hypothesis, we infected THP1 monocytes carrying an inducible PDGFRα with a triple fluorescent HCMV strain carrying fluorescent tags for immediate early (IE), early and late viral gene expression ^38^. At 16 hpi, prior to any viral genome replication, we sorted cells according to their IE expression to bright and dim populations (Fig. 7b). We found that the viral genome levels were >15-fold lower in dim compared to bright cells (Fig. 7c), indicating that the dim cells are the cells that initially received less viral genomes. The dim population was re-sorted at 5dpi to ensure there were no lytic cells which were not detected at 16hpi. We followed the two cell populations and found that at 7dpi the sorted IE-bright cells are indeed lytically infected, as they robustly expressed the late viral gene marker and produced infectious viral progeny while the IE-dim cells did not (Fig. 7d, S7a). Importantly, the isolated dim cells were capable of reactivation and release of infectious viral progeny upon differentiation at 7dpi (Fig. 7e, S7b), indicating they were latently infected. These data show that indeed cells that receive less viral genomes have a lower chance of becoming productively infected and are the ones in which latency is established.

**Fig. 7.**
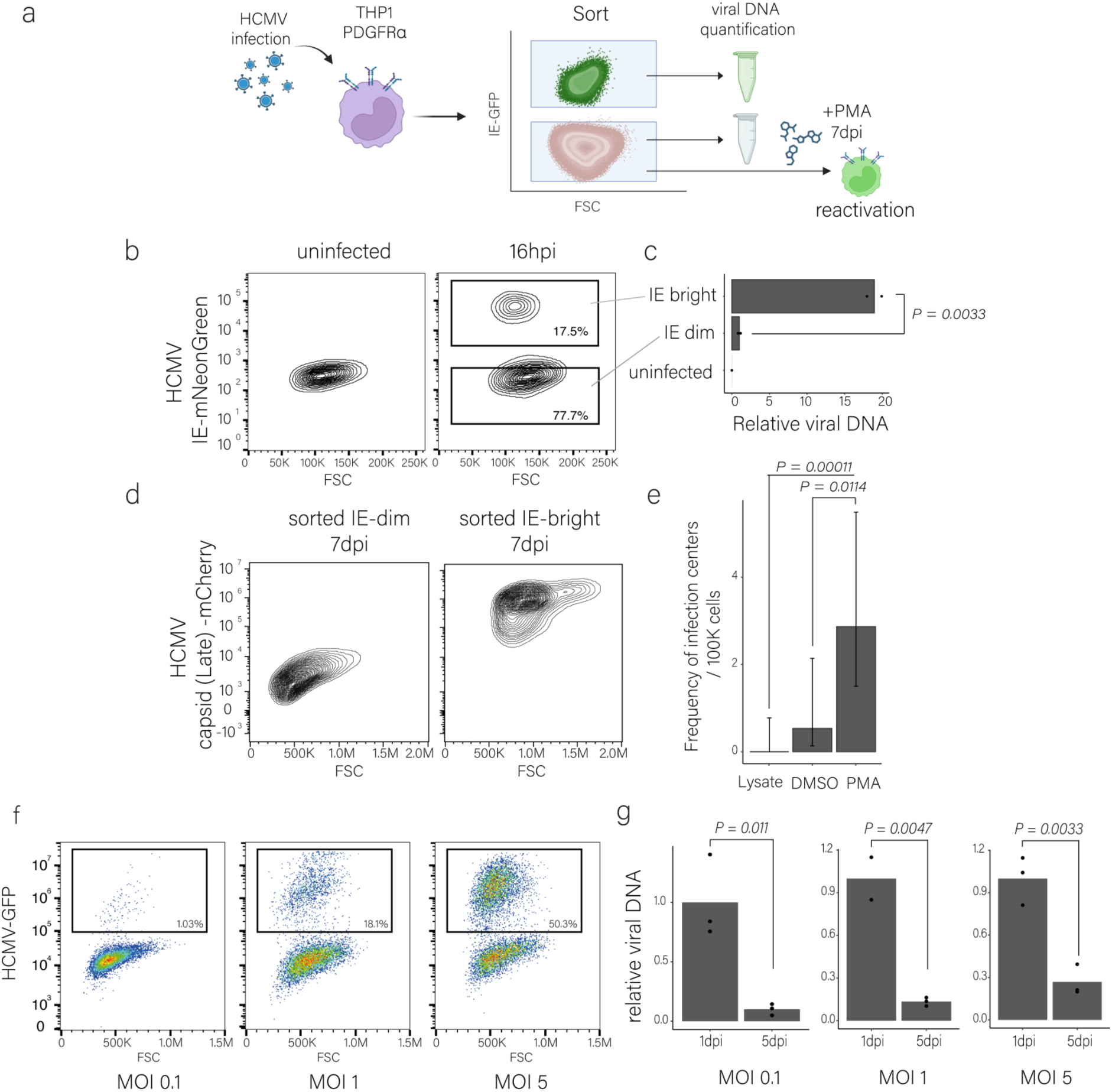
THP1-PDGFRa receiving less viral genomes establish latent infection. (a). Illustration of the experimental setup. (b). THP1 monocytes were induced to express PDGFRα and were infected with a triple fluorescent HCMV strain carrying fluorescent tags for immediate early (IE), early and late viral gene expression ^38^ (MOI=5). Cells were sorted at 16hpi based on mNeonGreen expression from the IE promoter to bright and dim populations. Gating strategy is shown in Fig. S1a. (c). Relative viral DNA levels in sorted bright and dim populations at 16hpi. Viral DNA levels were measured by real-time PCR and were normalized to a cellular genomic target. n=3. (d). Flow cytometry analysis at 7dpi of mCherry from the viral late gene promoter (UL48A) in HCMV-mNeonGreen (IE)-positive and negative sorted cells. Gating strategy is shown in Fig. S1a. (e). Reactivation levels of HCMV in sorted THP1–PDGFRα dim cells measured on fibroblasts as frequency of infectious centers calculated with ELDA software ^39,40^. Undifferentiated cells and cell lysates from 7dpi are shown as controls. (f). Flow cytometry analysis of THP1–PDGFRα infected with HCMV-GFP at MOIs of 0.1, 1 and 5. The dim population was sorted according to GFP expression at 5dpi. Gating strategy is shown in Fig. S1a. (g). Relative viral DNA levels from THP1–PDGFRα cells at 1dpi and from sorted dim cells at 5 dpi, measured from extracted DNA. Viral DNA levels were measured by real-time PCR and were normalized to a cellular genomic target. n=2-3.

To further substantiate these results, we infected THP1-PDGFRα with HCMV-GFP at different MOIs and sorted the dim cells at 5dpi (Fig. 7f). As expected, with increasing MOI the percentage of lytic cells increased. In all MOIs the viral load was significantly lower in the dim cells sorted at 5dpi compared to the viral load in all cells at the early stage of infection (Fig. 7g), indicating that these cells represent the cells that initially received less viral genomes and illustrating that the number of particles that infect a cell plays a major role in dictating infection outcome in monocytes. Across all MOIs tested, these dim cells retained the capacity of reactivation upon differentiation (Fig. S7c), indicating they were latently infected. These results propose a model by which the number of incoming viral genomes determines the likelihood of productive infection. When the number of viral genomes per cell falls below a critical threshold, the chances of viral genome repression increase, and latency can be established (Fig. 8).

**Fig. 8.**
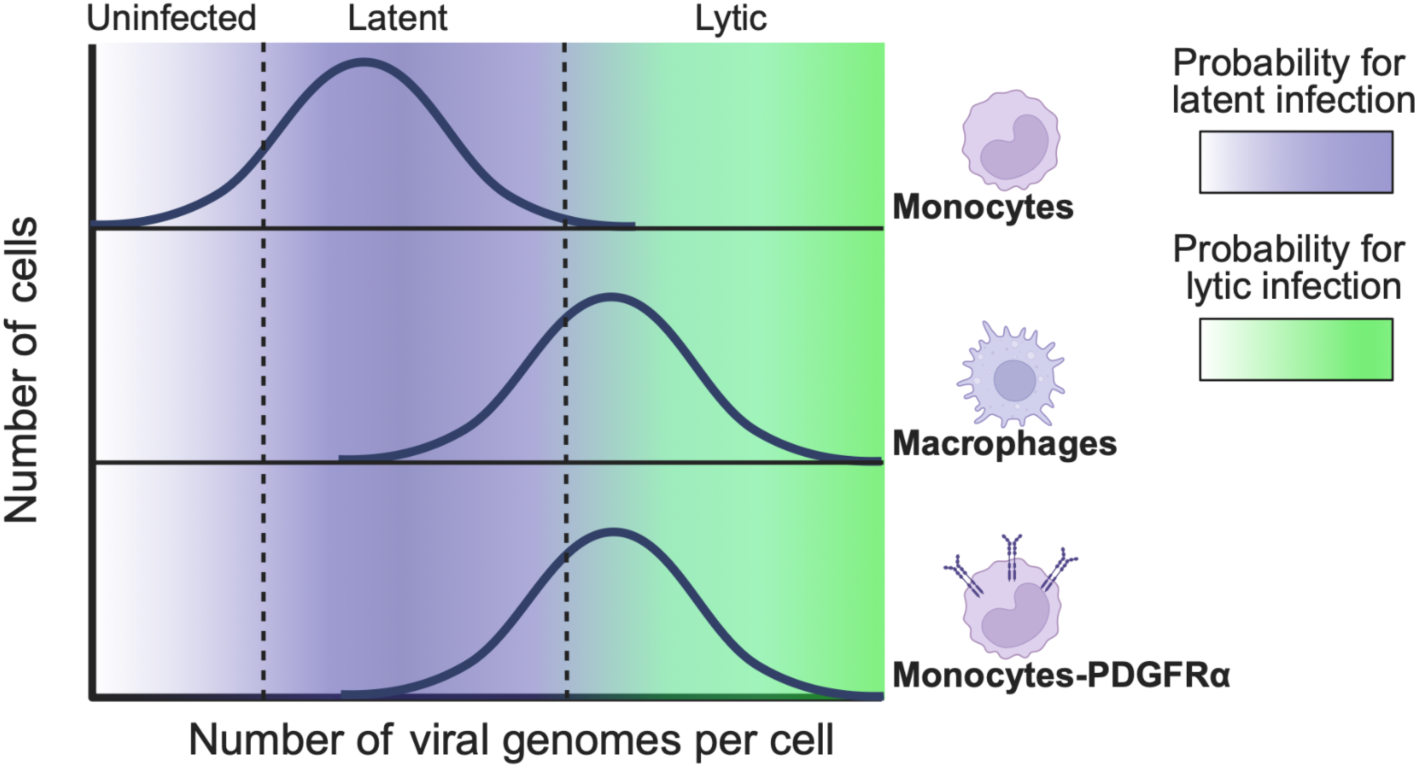
Model for the effect of viral entry on productive versus latent HCMV infection Schematic model illustrating the relationship between viral genome load and infection outcome. The histograms represent the distribution of viral genomes per cell within the population. The color gradients indicate the probability for latent infection (purple) or lytic infection (green), with dashed lines marking the approximate thresholds. Monocytes typically receive fewer viral genomes, favoring latency, compared to macrophages. Ectopic expression of an HCMV receptor, e.g. PDGFRα, in monocytes, increases the viral genome load driving them towards lytic infection.

Overall, these results demonstrate that inefficient entry of HCMV into monocytes is a major factor underlying the low levels of viral gene expression and, consequently, the ability to establish latency. Facilitating efficient entry of viral genomes enables productive replication even in monocytes, underscoring that entry is a critical determinant of infection outcome and latency establishment.

## Discussion

HCMV infection of monocytes results in a latent infection in which the virus is largely repressed and does not replicate, while following differentiation of these cells, they become permissive to productive infection and produce progeny.

Previous studies have shown that latent infection is characterized by low levels of viral transcripts ^41,42^. Using metabolic labeling, we show that viral gene transcription is very low in monocytes while substantial in macrophages. Chromatin regulation has been implicated as a major factor for viral repression during latency ^9,11^. Although some specific chromatin factors have been attributed to this specific repression ^43–45^, the differences between monocytes and their differentiated counterparts with regard to chromatin repression remain poorly defined. Our findings indicate that chromatin-based repression does not fully account for the inability of monocytes to support productive infection. Using HDAC inhibitors, chromatin repression can be relieved and indeed result in elevated productive infection in both monocytes and macrophages, but still productive infection in monocytes remains very limited.

We found a striking disparity in the abundance of viral genomes within the nuclei of infected monocytes compared to macrophages at early time points, in line with previous studies of monocyte infection ^46^, and notably, a significant difference in the quantity of viral capsids within these cells. This suggests HCMV entry efficiency as an unexplored barrier for productive infection in monocytes. Remarkably, ectopic expression of known HCMV entry receptors in monocytes facilitates productive infection, underscoring inefficient entry as a barrier in these cells. This means that although monocytes are capable of supporting productive infection, they do not reach a certain threshold of viral genome load required to establish productive infection due to less efficient viral entry to these cells (Fig. 8).

By performing unbiased transcriptome analyses, we interestingly found significant upregulation upon differentiation of several cell surface proteins which are linked to the entry of HCMV. One of these receptors was NRP2, which mediates HCMV entry into non-fibroblast cells, through the viral pentamer complex. Although its expression increased during differentiation, our results indicate that it does not play a major role in viral entry into macrophages, at least with the TB40 strain. ITGB3 and ITGB1 were both shown to play a role in HCMV entry ^12,36,47^ and depletion of these integrins showed that ITGB3, which is not expressed in monocytes, is required for HCMV entry into macrophages. However, overexpression of ITGB3 alone or with its canonical partner, ITGAV, was not sufficient to enable productive infection in monocytes, indicating the involvement of additional factors or post-translational modifications^48^ that are required for entry and likely absent or low in monocytes.

The notion that entry constitutes a major barrier for productive infection in monocytes suggests that the number of particles that infect a cell plays a major role in the probability of establishing productive versus latent infection. The finding that in infected monocytes, cells receiving less viral genomes fail to establish productive infection and instead enter latency further supports this conclusion. Therefore we suggest a model by which when the number of viral particles in a cell falls below a critical threshold, productive infection does not occur, likely due to the inability to prevent repression of the genome by the host, and instead latent infection may arise (Fig. 8). We show here that the composition of receptors on the cell surface affects entry in the case of monocyte to macrophage differentiation, however additional factors may affect viral genome entry, such as cell state, exposure to interferons, etc. Moreover, different cell types may have different thresholds for the amount of particles required for establishing productive infection. This may be affected for instance by the balance of activating versus repressing chromatin profiles. This raises the possibility that infection at low multiplicity, possibly in diverse cell types, may lead to a subset of cells harboring an insufficient amount of genomes or particles to support productive infection, these will likely be repressed and potentially still facilitate the establishment of latency.

Altogether, our findings identify inefficient entry as a critical barrier to productive infection in monocytes, underscoring an overlooked yet pivotal aspect of monocyte susceptibility to latent HCMV infection.

## Material and Methods

### Ethics statement

All fresh peripheral blood samples were obtained after approval of protocols by the Weizmann Institutional Review Board (IRB application 92-1) and following informed consent from the donors. Blood donors were not compensated.

### Cell culture and virus

Primary CD14+ monocytes were isolated from fresh venous blood, obtained from healthy donors, males and females, aged 25–45, using Lymphoprep density gradient (Stemcell Technologies) according to the manufacturer’s instructions, followed by magnetic cell sorting with CD14+ magnetic beads (Miltenyi Biotec) according to the manufacturer’s instructions. Monocytes were cultured in X-Vivo15 media (Lonza) supplemented with 2.25 mM L-glutamine at 37 °C in 5% CO2, at a concentration of 1–2 million cells per ml in non-stick tubes to avoid differentiation.

Where indicated, primary monocytes were differentiated immediately following isolation by culturing in RPMI with 20% heat-inactivated fetal bovine serum (FBS), 2 mM L-glutamine and 100 units ml−1 penicillin and streptomycin (Beit-Haemek) supplemented with 50 ng/ml PMA (Sigma) for 3 days. Culturing of monocyte derived macrophages was in RPMI with 20% heat-inactivated fetal bovine serum (FBS), 2 mM L-glutamine and 100 units ml−1 penicillin and streptomycin (Beit-Haemek).

293T cells (ATCC CRL-3216) and Primary human foreskin fibroblasts (ATCC CRL-1634) were maintained in DMEM with 10% FBS, 2 mM L-glutamine and 100 units ml−1 penicillin and streptomycin (Beit-Haemek).

THP1 cells, purchased from ATCC (TIB-202), and Kasumi-3 cells, purchased from ATCC (CRL-2725), were grown in RPMI media with 20% heat-inactivated FBS, 2 mM L-glutamine and 100 units/ml penicillin and streptomycin (Beit-Haemek). Differentiation of THP1 and Kasumi-3 cells was done by adding 50 ng/ml PMA for 3 days.

24h Before infection, cells were grown in 0.5% or 10% FBS For experiments in which the cells were collected at 3dpi or progeny, respectively, and supplemented with 10% FBS after 1h of infection.

In the experiments in which receptors were induced before infection, cells were treated with 1ug/ml doxycycline, with or without 1µM TrichostatinA (TSA, Sigma) for 24h before infection. The cells were validated for the receptor expression by surface staining and washed before HCMV infection.

TSA was added at the specified concentrations, at 5 hpi, DMSO was added as control.

Imatinib (Sigma) was added at the specified concentrations 1h before infection, DMSO was added as control.

IFNγ (500U/ml, peprotech) was added for 2 days to THP1 monocytes before surface staining of MHC-II.

The TB40E strain of HCMV, containing an SV40–GFP reporter, was described previously ^13^. For microscopy imaging, a previously reported TB40E strain containing GFP fused to a tegument protein (UL32-GFP) was used ^30^. For the reactivation experiment presented in Fig. 7b-e, a previously described TB40 strain expressing triple fluorescence was used ^38^.

Virus propagation was done by adenofection of a bacterial artificial chromosome of the viral genome into fibroblasts as described previously (Elbasani et al., 2014). When most of the cells in the culture died, supernatant was collected and cleared from cell debris by centrifugation.

### Infection procedures

Infection was performed by centrifugation at 800g for 1h in 24-well or 12-well plates with the virus added at a multiplicity of infection (MOI) = 5, unless stated differently, followed by washing and supplementing with fresh media. Notably, because this MOI is based on quantification of infectious particles in fibroblasts it is effectively lower in monocytic cells.

For progeny assay, at 8 or 10 dpi. The supernatant was cleared from cell debris by centrifugation and transferred to fibroblasts. Infected fibroblasts were counted 2-3 days later.

For Reactivation assays described in figures S1b and S7c, GFP dim cells were sorted at the indicated time point, as specified (in monocytes with no PDGFRα expression we validated there are no GFP-bright cells) and at 7dpi were treated with PMA (50ng/ml) or with DMSO as control. HCMV-positive cells were counted on a fluorescent microscope. Reactivation assay using Extreme Limiting Dilution Assay (ELDA), described in Fig. 7e, was performed as previously described ^39^at 7 dpi, cells were seeded with PMA or with DMSO, as control, in serial dilutions ranging from 20,000 cells to 625 cells per well in 96 well plate, with 8 replicates per dilution. The equivalent number of cells were lysed and seeded to control for infectious virus in the initial cells. After 2 days, 10,000 fibroblasts were added to each well to allow detection of infectious virus. At 24 dpi, plaques were counted, and reactivation levels were quantified using ^40^.

### Flow cytometry and sorting

Adherent cells were harvested by washing with PBS and in 0.5 mM EDTA before scraping. Cells were analyzed on a BD Accuri C6 or CytoFLEX (Beckman Coulter) and sorted on a BD FACS AriaIII using FACSDiva software. All analyses and figures were done with FlowJo. All histograms were plotted with modal normalization.

### Next-generation sequencing

RNA-seq library preparation was performed as described previously ^49^. Cells were collected with Tri-Reagent (Sigma-Aldrich). Total RNA was extracted according to the manufacturer’s instructions, and poly-A selection was performed using Dynabeads mRNA DIRECT Purification Kit (Invitrogen). The mRNA samples were subjected to DNase I treatment and 3ʹ dephosphorylation using FastAP Thermosensitive Alkaline Phosphatase (Thermo Scientific) and T4 polynucleotide kinase (New England Biolabs) followed by 3ʹ adaptor ligation using T4 ligase (New England Biolabs). The ligated products were used for reverse transcription with SSIII (Invitrogen) for first-strand cDNA synthesis. The cDNA products were 3ʹ-ligated with a second adaptor using T4 ligase and amplified for 8 cycles in a PCR for final library products of 200–300 bp. Raw sequences were first trimmed at their 3’ end, removing the Illumina adapter and poly(A) tail. Alignment was performed using Bowtie 1^50^ (allowing up to two mismatches) and reads were aligned to the human (hg19). Reads aligned to ribosomal RNA were removed. Reads that were not aligned to the genome were then aligned to the transcriptome.

For sequencing the genome of the HCMV-GFP virus used for infection, DNA was extracted from infected THP1-PDGFRα cells using blood kit (QIAGENE). The sequencing library was prepared using NEBNext® DNA Library Prep Kit according to the manufacturer’s instructions. Reads were aligned to the TB40 reference genome (EF999921) using STAR. The resulting wiggle file was inspected manually, and no mutations or deletions were detected in the genes encoding the viral entry receptors.

### Differential expression and enrichment analysis

Differential expression analysis on RNA-seq data was performed with DESeq2 (v.1.22.2) using default parameters, with the number of reads in each of the samples as an input.

The log2(fold change) values from the DE on the RNA-seq was used for enrichment analysis using GSEA (v.4.1). For the analysis, only genes with a minimum of ten reads were included.

Gene lists of cellular receptors^28^, chromatinization^51^ and ISGs^52^ are based on the referred papers. Monocyte to macrophage differentiation gene list is based on the list of genes in the GO term regulation of macrophage differentiation on GO:0030225.

### RNA labeling for SLAM-seq and analysis

For metabolic RNA labeling, 4sU (T4509, Sigma) was added at a final concentration of 200 μM to infected cells at 3hpi. Cells were collected with Tri-reagent at 2 and 3 h after adding the 4sU (corresponding to 5 and 6 hpi). RNA was extracted under reducing conditions and treated with iodoacetamide (A3221, Sigma) as previously described ^16^. RNA-seq libraries were prepared and sequenced as described in the ‘Next generation sequencing’ section.

Alignment of SLAM-seq reads was performed using STAR, with parameters that were previously described ^53^. First, reads were aligned to a reference containing human rRNA and tRNA, and all reads that were successfully aligned were filtered out. The remaining reads were aligned to a reference of the human (hg19) and the TB40 (EF999921). In one analysis, the virus was analyzed as one transcript, and in a second analysis, all viral genes were analyzed. Output.bam files from STAR were used as input for the GRAND-SLAM analysis ^54^ with default parameters and with trimming of 5 nucleotides in the 5ʹ and 3ʹends of each read. Each one of the runs also included an unlabeled sample (no 4sU) that was used for estimating the linear model of the background mutations. The estimated ratio of newly synthesized out of total molecules for each viral and host gene were used for the presented analyses.

### 3D immunofluorescence viral DNA FISH

Differentiated monocytes (THP1 and CD14+) were seeded on 22X22 coverslips in a 6-well plate. Suspension monocytes were concentrated to 1M cells/150μl and seeded on 22X22 coverslips for 1 hour, followed by centrifuging for 10 min in 800g to deposit suspension cells onto the coverslips. FISH was done as previously described^55^. Cells were washed twice with PBS and fixed in 4% paraformaldehyde in PBS for 10 minutes. Then, cells were permeabilized in 0.5% triton/PBS for 15 minutes and rinsed in PBS. Samples were incubated in 20% glycerol/PBS at 4°C overnight and frozen five times in liquid nitrogen. Cells were washed three times in 0.05% triton/PBS, rinsed with PBS followed and with DDW. Cells were then incubated in 0.1M HCL for 15 minutes and then with 0.002% pepsin (Sigma, P6887)/0.01M HCl at 37°C for 90 seconds for monocytes or 75 seconds for macrophages, followed by inactivation in 50mM Mgcl2/PBS. Cells were then fixed in 1% paraformaldehyde/PBS for 1 minute and washed with PBS and with 2X SSC (Promega, V4261), followed by Incubation in 50%formamide/2X SSC (pH7-7.5) for 1 hour.

Probe was prepared from a TB40 SV40-GFP BAC DNA using a Nick Translation Mix (Roche, 11745808910, Tetramethyl-Rhodamine-5-dUTP, Roche 11534378910) according to the manufacturer instructions, and prepared for hybridization by combining 0.5mg of each labeled probe and 5mg cot-1 (Invitrogen, 15279-011) in 4.5μl deionized formamide (Sigma, F9037) and 4.5μl 4XSSC/20% dextran sulfate.

Following denaturation at 76°C, hybridization was performed at 37°C for 3 days in a humid chamber. After hybridization, the coverslips were washed in 50% formamide/2X SSC (pH 7-7.5) at 37°C, in 0.5XSSC at 60°C and in 4XSSC/0.2%. Coverslips were mounted on slides with Prolong gold (Invitrogen, P36930) containing DAPI.

### Immunofluorescence

For HCMV replication compartment detection, cells were plated on i-bidi slides, fixed in 4% paraformaldehyde for 15 min, washed in PBS, permeabilized with 0.1% Triton X-100 in PBS for 10 min and then blocked with 10% goat serum in PBS for 30 min. Immunostaining was performed for the detection of mouse anti-UL44 (CA006-100, Virusys) with 2% goat serum diluted in PBS. Cells were washed 3 times with PBS and labeled with goat anti-mouse–Alexa Fluor 647 (Thermo Fisher) secondary antibody and DAPI (4ʹ,6-diamidino-2-phenylindole) diluted 1:500 in PBS for 1 hour at room temperature, followed by 3 PBS washes.

Detection of the HCMV tegument protein (pp150, UL32) in the cytosol was performed by infecting the cells with the TB40-UL32-EGFP ^30^ for one hour, followed by three PBS washes and 15 minutes of fixation with 4% paraformaldehyde. Samples were mounted with DAPI for nuclear staining and Phalloidin, which binds F-actin, to define cell borders.

### Microscopy and Image Analysis

DNA-FISH and HCMV capsid images were taken using Leica TCS SP8 CLSM. DNA-FISH images were analyzed using Imaris 10 software. Image files were anonymized by renaming and mixing to ensure blinded analysis. Positive signals were manually counted for each cell.

Infected cells with fluorescently labeled HCMV tegument protein (UL32-GFP) were visualized and analyzed using ImageJ (FIJI, NIH). Automated segmentation of individual virions was performed using the StarDist plugin^56^, while cell boundaries were manually delineated to define individual cell regions of interest (ROIs). Virions located within each cell ROI were assigned accordingly, enabling quantification of viral entry at single-cell resolution.

Images of HCMV replication compartment, bright field images and images of GFP positive infected cells were acquired on an AxioObserver Z1 wide-field microscope and analyzed using ImageJ.

### Cell Surface Staining

Cells were washed three times with PBS and blocked for 15 min in 2% human serum. After blocking, cell staining was done using the following conjugated antibodies: Allophycocyanin (APC)-conjugated mouse IgG2a anti-human Neuropilin-2 (R&D systems, catalog no. FAB22151A) with allophycocyanin (APC)-conjugated mouse IgG2a control (R&D systems, catalog no. IC003A). Phycoerythrin (PE)-conjugated Mouse IgG2a anti-human PDGFRα (BD, catalog no. 556002) with phycoerythrin (PE)-conjugated Mouse IgG2a control (BD, catalog no. 555574). Alexa fluor 647 conjugated Mouse IgG1 anti-human CD61 (ITGB3) (BLG, catalog no. 336407) and Alexa fluor 647 conjugated Mouse IgG1 control (BLG, catalog no. 400130). FITC-conjugated Mouse IgG1 anti-human CD29 (ITGB1) (Santa cruz catalog no. MEM-101A) and FITC-conjugated Mouse IgG1 control (Santa Cruz, catalog no. sc-2339). APC-conjugated Rat IgG2b, k anti-human CD11b (BioLegend catalog no. 101211). APC-conjugated Rat IgG1, k anti-human CD115 (BioLegend catalog no. 347305) and PE-conjugated Mouse IgG2b anti-human CD11c (BD catalog no. 333149). APC-conjugated Anti-HLA-DR mouse IgG2aκ (130-113-963).

Antibody incubation was done for 30 minutes at a 1:200 dilution. After staining, cells were washed twice with PBS and analyzed by flow cytometry.

### Plasmid construction lentiviral transduction

PDGFRα was cloned into pLex_TRC206 plasmid (Straussman et al., 2012) under EF-1α promoter and blasticidin selection. PDGFRα was amplified from cDNA (Supplementary Table 5) and cloned into pLex by FastCloning^57^.

For CRISPR knockout plasmids of NRP2, ITGB3 and ITGB1, we cloned two gRNAs into the lentiCRISPRv2-2guide plasmid using restriction-free cloning as previously described^58^ (Supplementary Table 5). Trip10 was used as a knock out control as described in ^58^.

To generate inducible expression plasmids of THBD, NRP2, ITGB3 and ITGAV, genes were ordered from TWIST. PDGFRa was amplified from the pLEX-PDGFRα plasmid. The genes were cloned into pLVX-Puro-TetONE-SARS-CoV-2-nsp1-2XStrep (kind gift from N. Krogan, UCSF) in place of the SARS-CoV-2-nsp1-2XStrep cassette using linearization with BamHI and EcoRI (neb). The genes were amplified with primers containing flanking regions homologous to the vector (Supplementary Table 5). The amplified PCR fragments were cleaned using a gel extraction kit (QIAGEN) according to the manufacturer’s protocol and were cloned into the vectors using a Gibson assembly reaction (neb). Inducible mCherry, cloned on the same backbone was previously described ^8^ and was used as control in all experiments with these inducible expression plasmids.

Lentiviral particles were generated by cotransfection of the expression constructs and second-generation packaging plasmids (psPAX2, Addgene, catalog no. 12260 and pMD2.G, Addgene, catalog no. 12259), using jetPEI DNA transfection reagent (Polyplus transfection) into 293T cells, according to the manufacturer’s instructions. At 60 h post-transfection, supernatants were collected and filtered through a 0.45-μm polyvinylidene difluoride filter (Millex). THP1 cells were transduced with lentiviral particles by centrifugation at 800g for 1h in 24-well or 12-well plates. 2 days post transfection the cells were transferred to selection media (blasticidin, 10 μg/ ml for 5 days or puromycin, 1.75 μg/ml for 4 days). Blasticidin and puromycin were removed and cells were recovered for at least two days before subsequent processing.

### CRISPR and siRNA-Mediated Knockdown

CRISPR-mediated knockout of *NRP2* and *ITGB1* was performed in bulk populations. For *ITGB3*, knockout was performed by CRISPR, followed by clonal selection of a single-cell-derived population and confirmation of mutations in both alleles by sequencing. In parallel, a control knockout of *TRIP10* was generated in a bulk population for comparison.

siRNA-mediated knockdown of *NRP2* and *ITGB3* was conducted using ONTARGETplus SMARTpool reagents (Dharmacon; L-017721-00-0005 for *NRP2* and L-004124-00-0005 for *ITGB3*), with a non-targeting control (D-001810-10-05). siRNAs were first diluted to 5 µM in siRNA buffer, then further diluted to 0.25 µM in Opti-MEM to prepare Solution I. In parallel, 4 µL of Lipofectamine RNAiMAX (Thermo Fisher) was diluted in 96 µL Opti-MEM to generate Solution II. Solutions I and II mixed together and were incubated for 20 minutes at room temperature.

The transfection mixture was added to THP1-derived macrophages at one day post-differentiation, in RPMI medium supplemented with 50 ng/mL PMA, to a final volume of 1 mL per well in a 12-well plate. Cells were incubated with siRNA complexes for 48 hours prior to HCMV infection.

### Quantitative real-time PCR analysis

Total RNA was extracted using Direct-zol RNA Miniprep Kit (Zymo Research) following the manufacturer’s instructions. cDNA was prepared using qScript FLEX cDNA Synthesis Kit (Quanta Biosciences) following the manufacturer’s instructions. Real-time PCR was performed using SYBR Green PCR master-mix (ABI) on the QuantStudio 12K Flex (ABI). Amplification of NRP2 and ITGB3 was normalized to the host gene ANAX5 (primers detailed in table S5).

For whole cell analysis, total DNA was extracted using QIAamp DNA Blood kit (Qiagen) according to the manufacturer’s instructions. For virion DNA extraction, samples were first treated with DNAseI to remove viral DNA not enclosed in intact virions (PerfeCTa, 95150), following the manufacturer’s instructions. 0.4 mg/ml proteinase K (Invitrogen, 25530049) was added and the samples were incubated for 1 h at 60 °C and then 10 min at 95 °C.

Amplification of the viral gene UL44 was normalized to the host gene B2M (primer detailed in table S5).

**Table S5.**
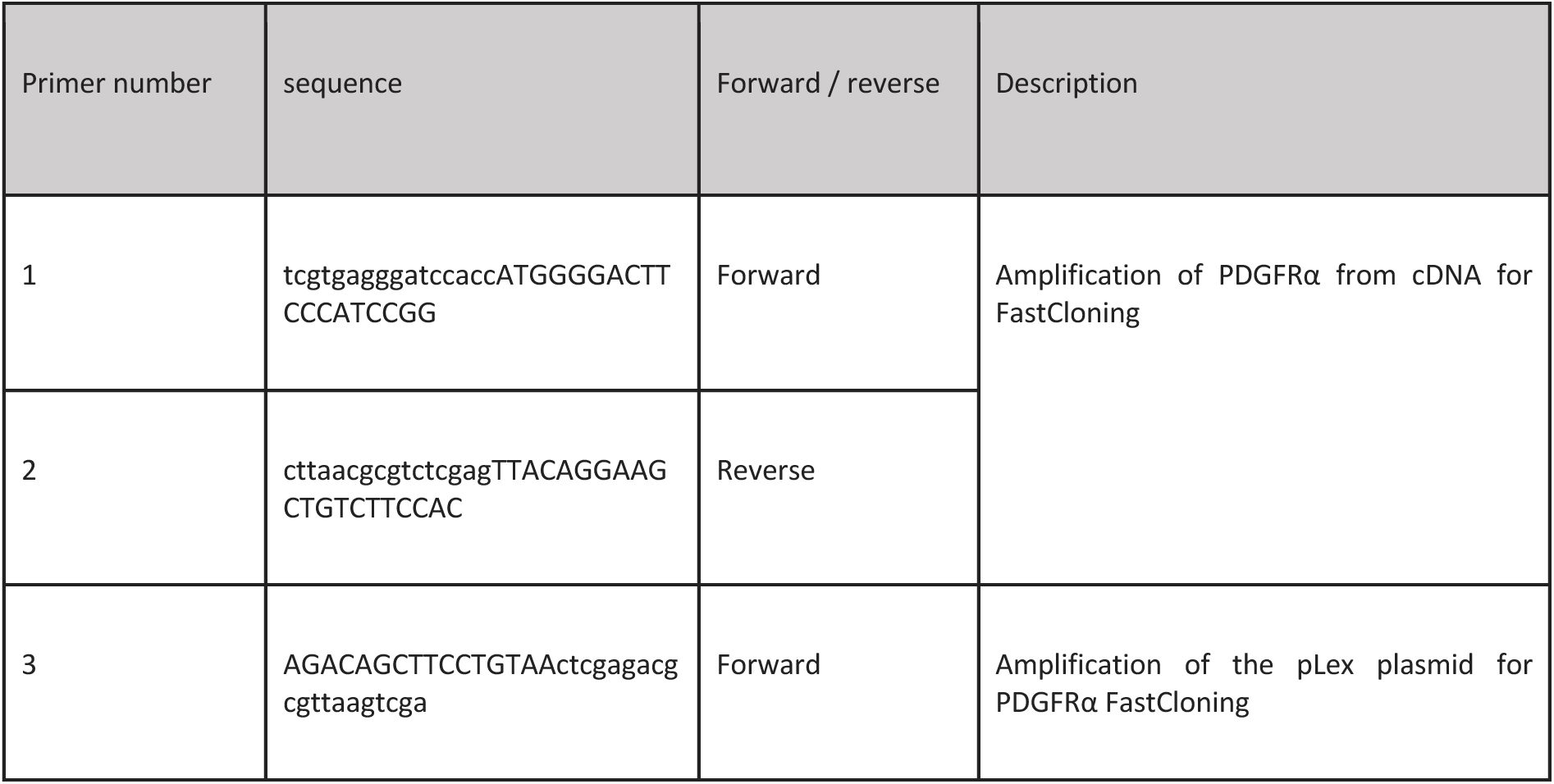
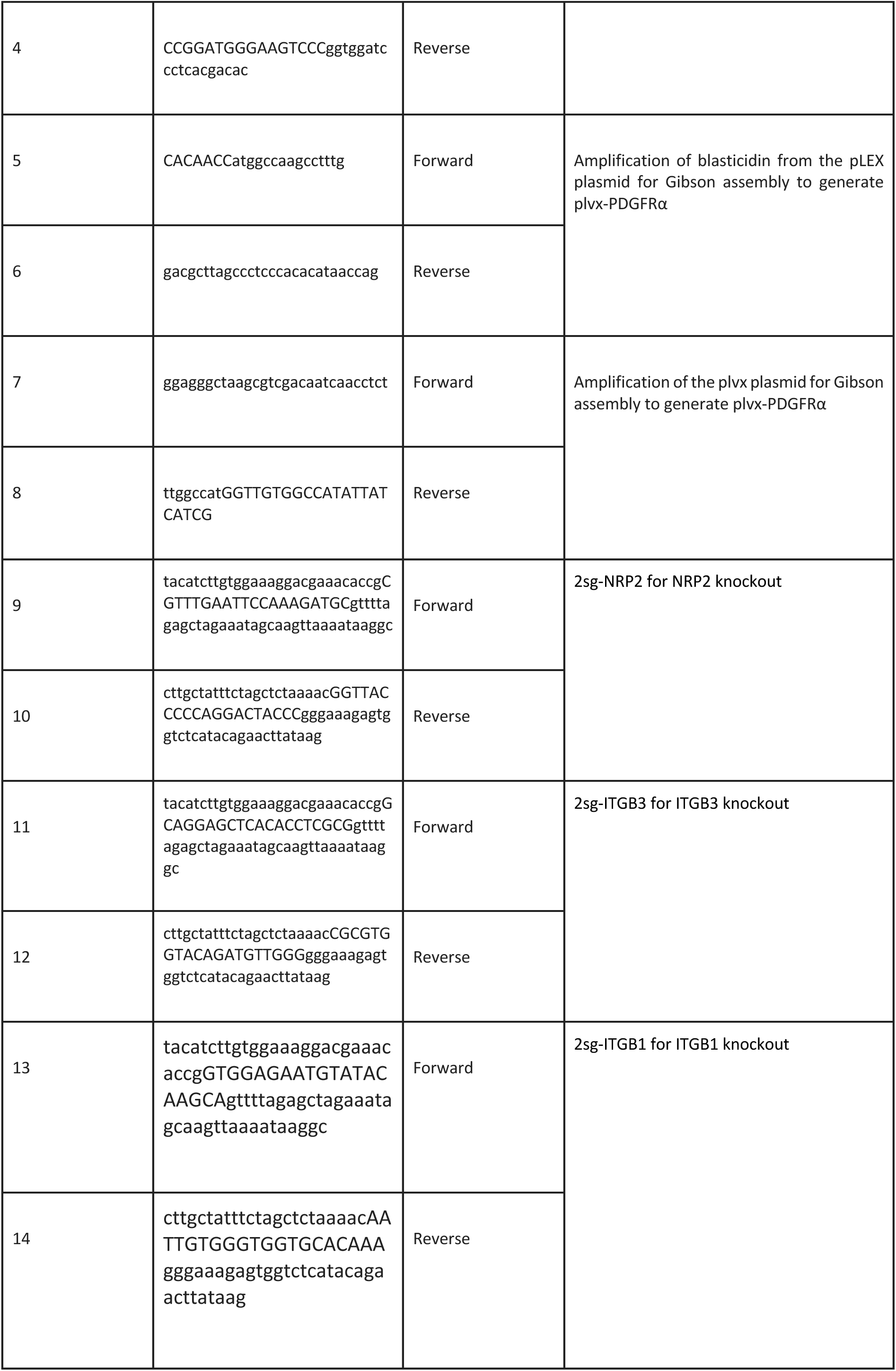
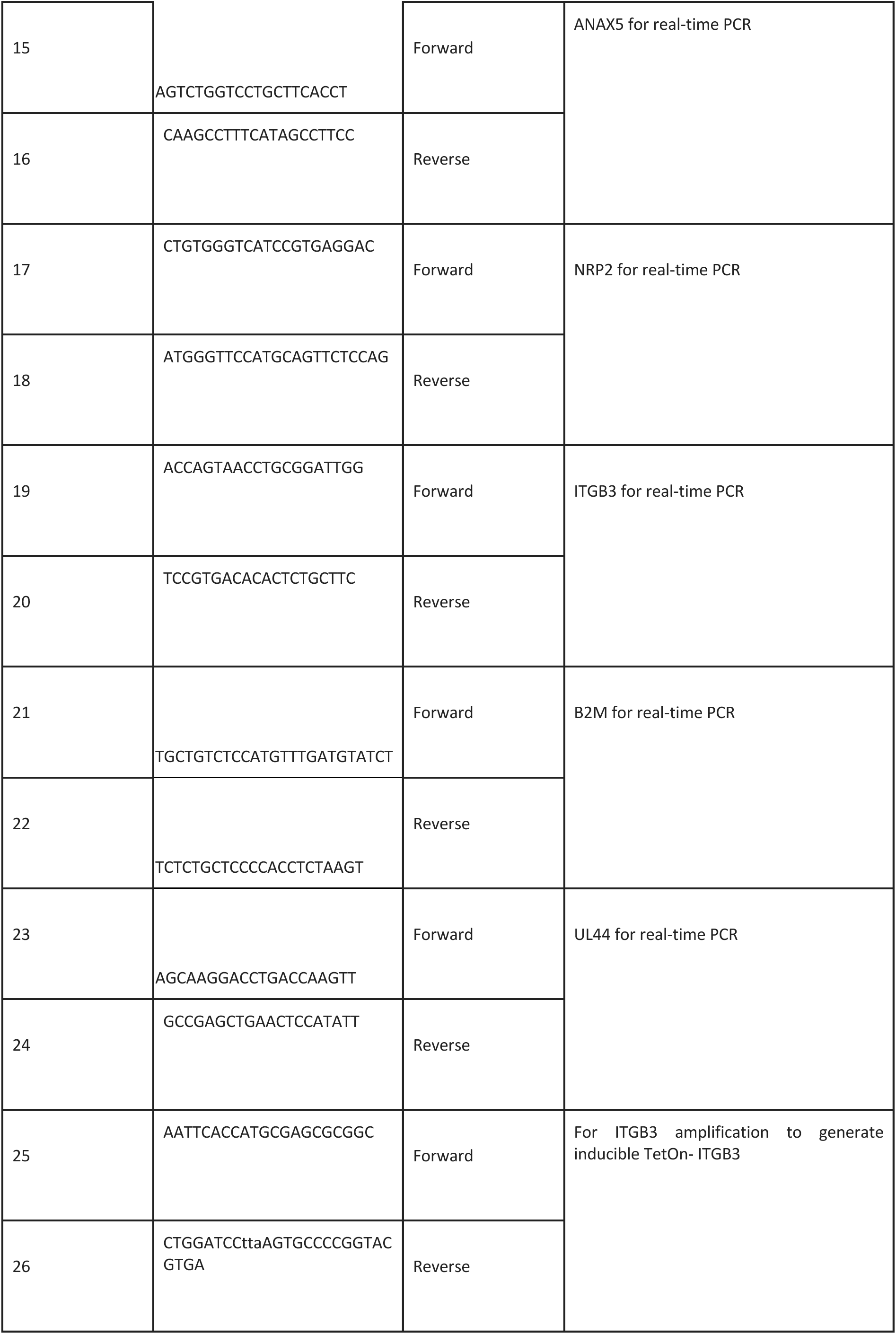
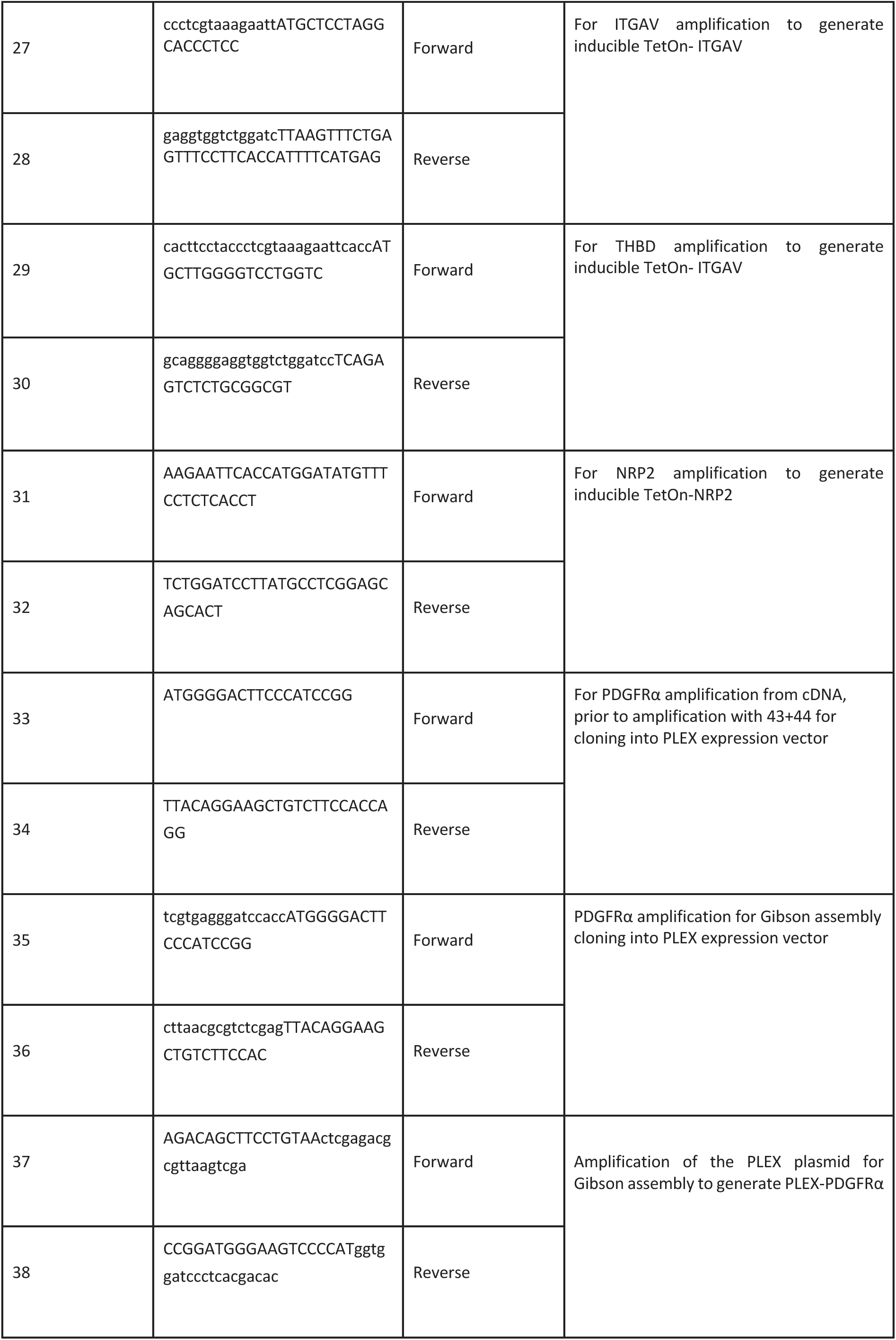
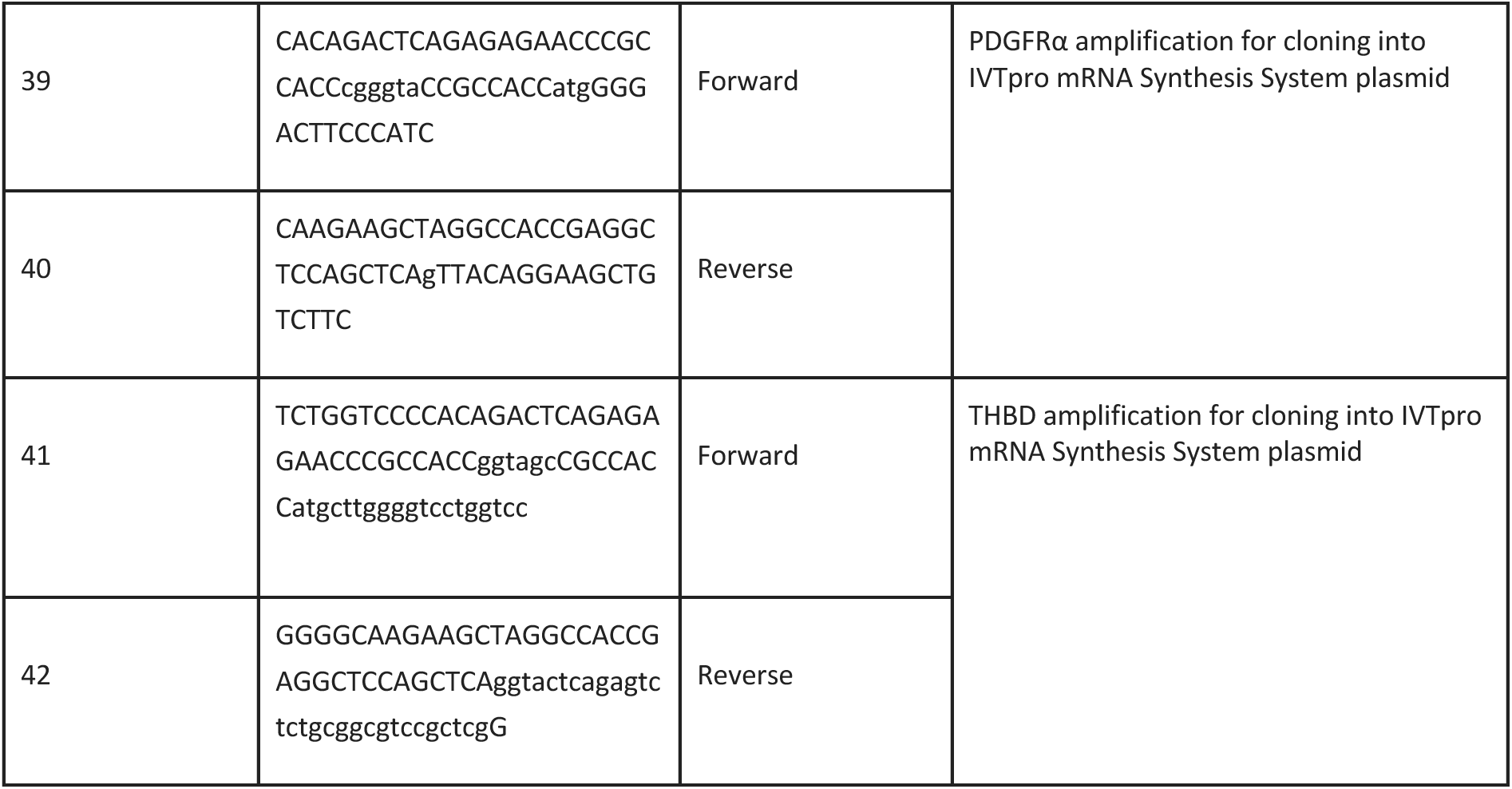
primers list.

### RNA In vitro transcription

PDGFRα and THBD were amplified using the primers detailed in table S5 and cloned into the DNA template plasmid (Takara IVTpro mRNA Synthesis System) using Gibson reaction and linearized with HindIII-HF (NEB) for following *in vitro* transcription reaction. IVT RNA was produced using the CleanScribe™ RNA Polymerase (E-0107, TriLink Biotechnologies) according to the manufacturer’s instructions. UTP was substituted with N1-Methylpseudouridine-5’-Triphosphate (N-1081, TriLink) and the RNA was co-transcriptionally capped with CleanCap Reagent AG (N-7113, TriLink). The RNA was treated with DNAseI (ON-109, Hongene), precipitated with LiCl and reconstituted in ddH2O. Primary monocytes were transfected with the produced RNA with jetMESSENGER® mRNA transfection reagent (Polyplus), together with 4uM Ruxolitinib for 12h before HCMV infection.

### Western blot

For western blot analysis cells were lysed in RIPA buffer (150 mM NaCl,1% NP-40, 0.5% sodium deoxycholate, 0.1%SDS, 50 mM Tris-HCl pH 7.4, and 1×EDTA-free protease inhibitor cocktail). Lysates were cleared by centrifugation and supplemented with Sample Buffer. Proteins were separated by SDS-PAGE electrophoresis, transferred to nitro-cellulose membranes (0.25 mm, ThermoFisherScientific), and detected using an infrared fluorescent antibody detection system (LI-COR) using the antibodies PDGF Receptor α Antibody (#3164, Cell Signaling Technology) and GAPDH (2118S, Cell Signaling Technology) as loading control.

### Mass spectrometry

The samples were lysed and digested with trypsin using the S-trap method. The resulting peptides were analyzed using nanoflow liquid chromatography (nanoAcquity) coupled to high-resolution, high mass accuracy mass spectrometry (TIMS-TOF Pro). Each sample was analyzed on the instrument separately in a random order in DIA mode. The resulting data was analysed using Spectronaut software package using the Direct-DIA workflow.

## Data availability

All next-generation sequencing data files have been deposited in Gene Expression Omnibus under accession number GSE280650.

## Acknowledgments

We thank Dr. Orly Laufman, Prof. Yossef Shaul and the members of the Stern-Ginossar lab for the critical reading of the manuscript. We thank Dr. Dor Simkin, Dr. Avner Leshem and Tatiana Smirnova for technical assistance. All optical imaging acquired at the de Picciotto Cancer Cell Observatory In memory of Wolfgang and Ruth Lesser of the Moross Integrated Cancer Center in the Department of Life Science Core Facilities, Weizmann Institute of Science. We would like to thank the Weizmann flow cytometry and microscopy unit for technical assistance. We’d like to thank David Morgenstern from the protein profile unit at the G-INCPM, the Weizmann Institute, for his assistance with proteomics analysis.

This study was supported by a European Research Council consolidator grant (CoG-2019-864012) and an Israel Science Foundation grant to N.S.-G. (2507/23).

## Author contributions

Y.K., N.S.-G. and M.S. conceived and designed the project. Y. K., T.A., A.W. and M.S. performed the experiments. T.F. helped analyze the HCMV particle images. Y.F. offered valuable guidance throughout the research. Y.K., A.N., N.S-G and M.S. analyzed and interpreted the data. Y.K., N.S.-G. and M.S. wrote the manuscript with input from all the authors.

---

**Fig. S1.**
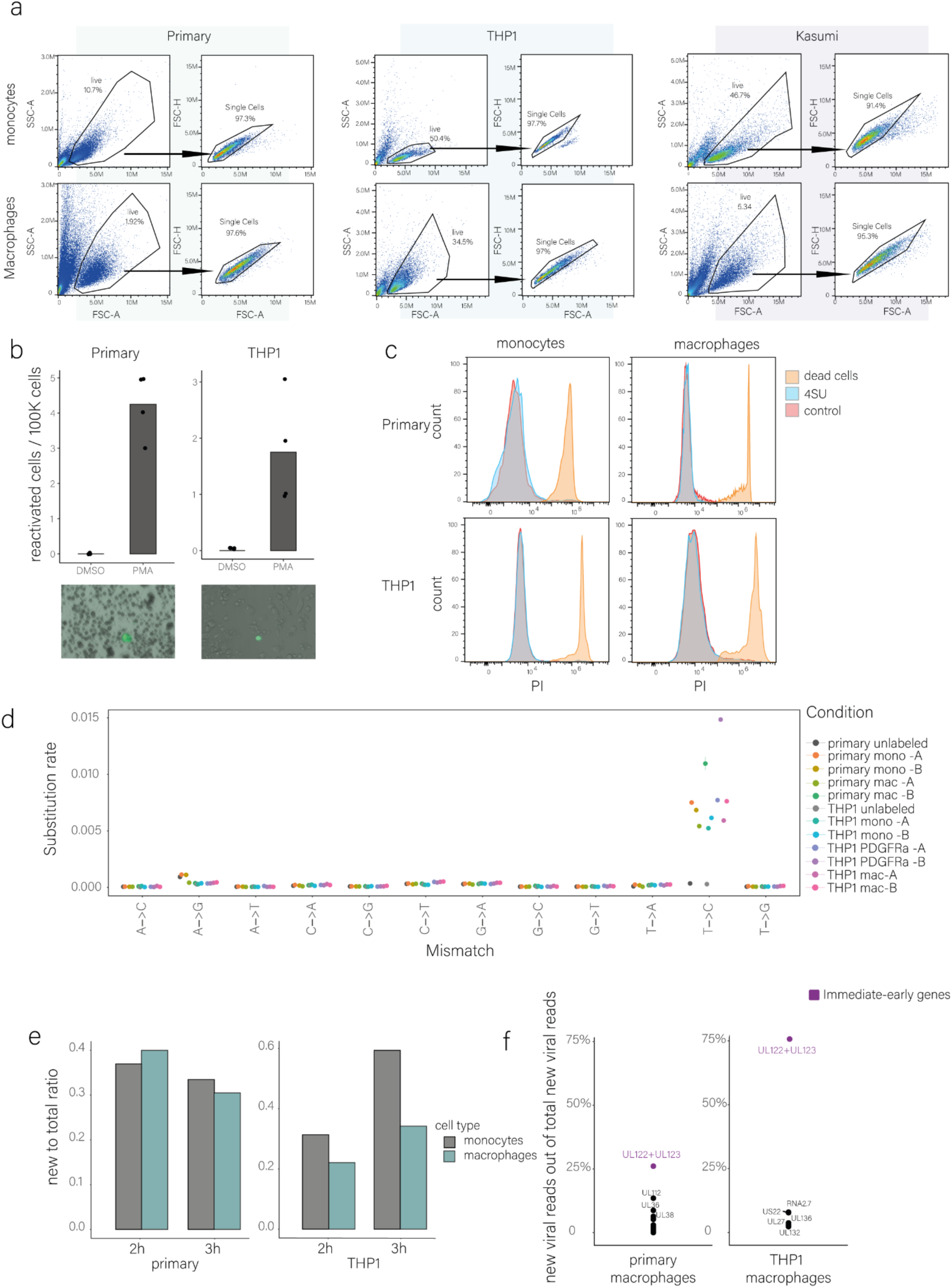
(a). Gating strategy used to determine the single-cell population for all flow cytometry analyses of monocytes and macrophages. (b). Primary and THP1 monocytes were infected with HCMV-GFP (MOI=5). At 7 dpi, cells were treated with either DMSO or PMA. 3 days post PMA treatment, amount of GFP positive monocytes were counted. (c). Survival assay of primary and THP-1 monocytes and macrophages. Cells were treated with 4-thiouridine (4sU) or water as control for 3 hours, followed by propidium iodide (PI) staining and flow cytometry analysis to assess cell viability. PI-positive cells represent non-viable populations, boiled cells were used to mark the dead cell population. (d). Rates of nucleotide substitutions demonstrate efficient conversion rates in 4sU-treated samples compared to unlabeled cells (no 4sU, gray dots) (e). Proportion of new-to-total RNA of cellular transcripts in infected primary and THP1 monocytes and macrophages. Infected cells were labeled with 4sU at 3 hpi and collected for RNAseq after two (left bars) or three (adjacent right bars) hours post labeling. (f). Percentage of new viral gene reads out of total new viral reads in infected primary and THP1 macrophages at three hours post labeling (6 hpi).

**Fig. S2.**
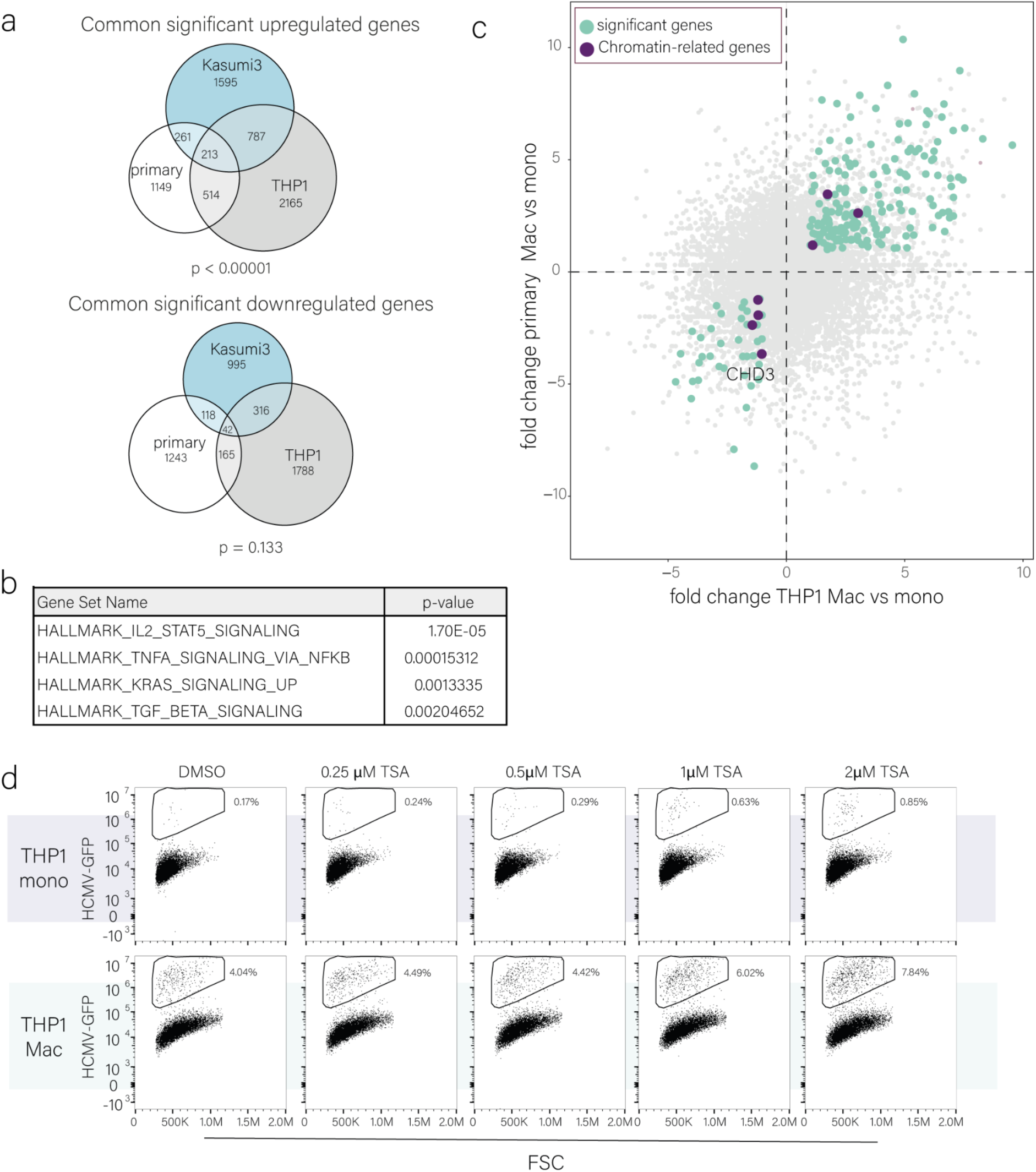
(a). Venn diagram summarizing the amount of significant upregulated and downregulated genes in all three cell types. Statistics on the common upregulated and downregulated genes were performed using hypergeometric test. (b). Hallmark pathway enrichment analysis shows four significantly enriched pathways in the 213 common significant upregulated genes. Analysis was performed using a hypergeometric test. (c). Scatterplot of the fold change (FC) from RNA-seq data between primary monocytes and macrophages, relative to the fold change between THP1 monocytes and macrophages. Light blue dots represent significantly upregulated genes in all three cell types. Dark purple dots represent chromatin-related genes that are significantly changing upon differentiation in all three cell types (p = 0.885 for the downregulated and p = 0.999 for the upregulated genes using a hypergeometric test), based on the EpiFactors database ^51^. (d). Flow cytometry analysis of infected THP1 monocytes and macrophages, treated with TSA or DMSO as control at 5 hpi. Analysis was performed at 3 dpi. The gate marks the productive, GFP-bright cell population. Gating strategy is shown in Fig. S1a.

**Fig. S3.**
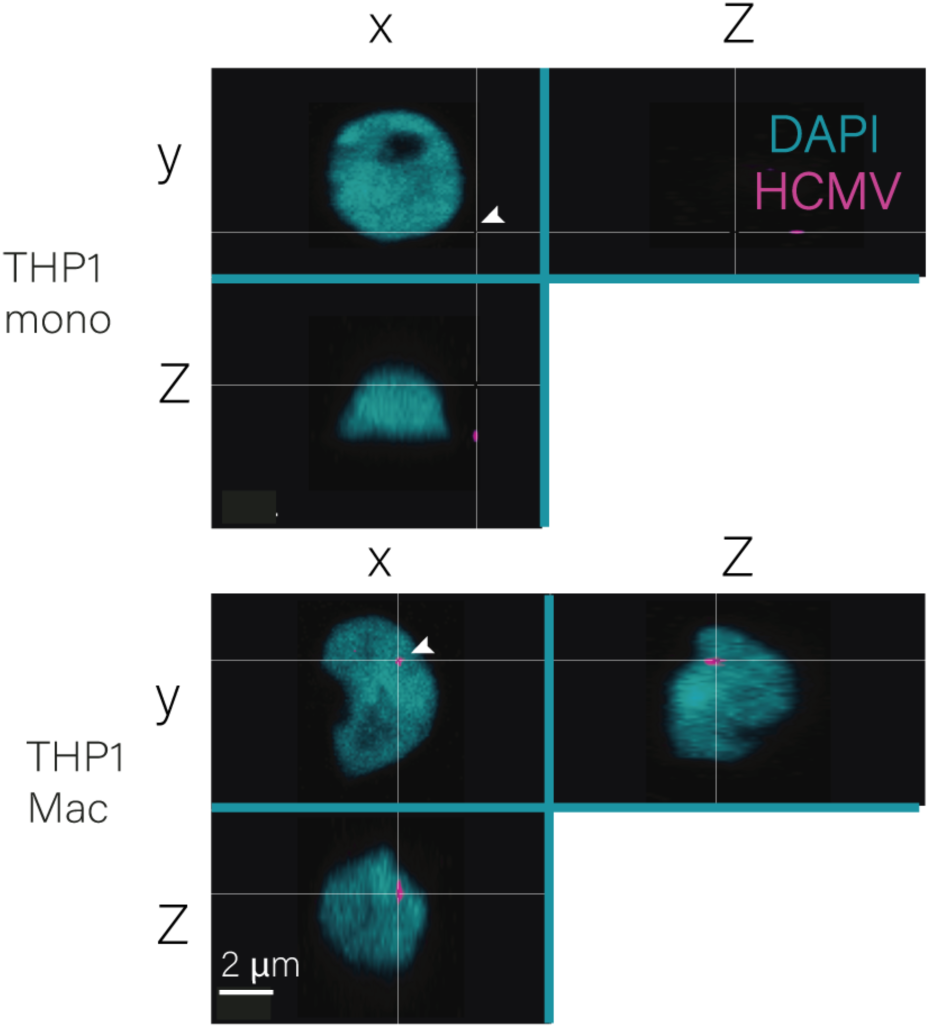
3D images of infected THP1 monocyte and macrophage nuclei at 12 hpi. The HCMV genome was probed using 3D DNA-FISH

**Fig. S4.**
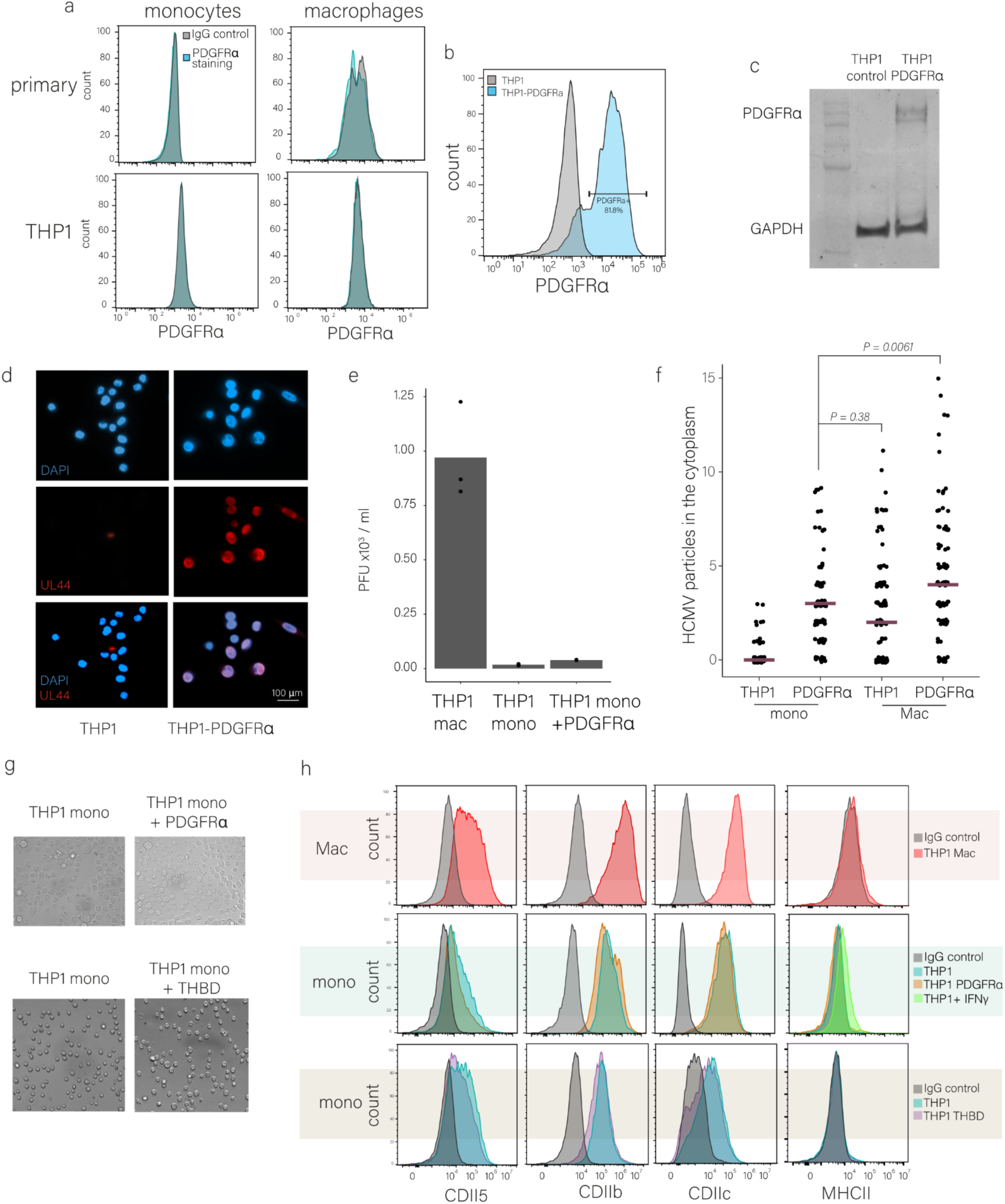
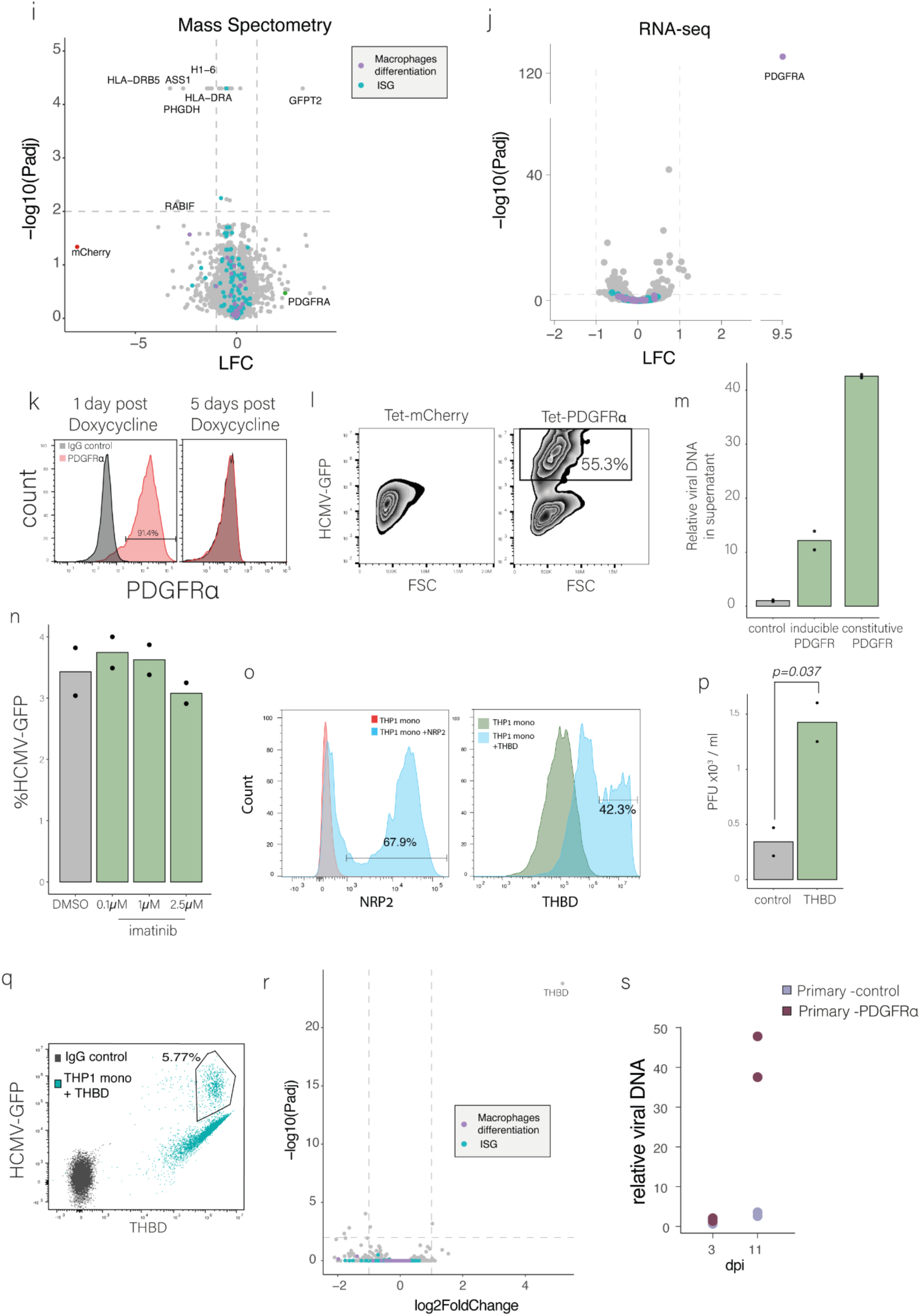
(a). Flow cytometry analysis of cell surface staining of PDGFRα versus IgG control in primary and THP1 monocytes and macrophages. Gating strategy is shown in Fig. S1a. (b). Flow cytometry analysis of PDGFRα surface expression on THP1 and THP1–PDGFRα monocytes. Gating strategy is shown in Fig. S1a. (c). Western blot analysis of THP1 monocytes overexpressing control or PDGFRα. GAPDH was used as loading control. (d). THP1 and THP1-PDGFRα were infected with HCMV-GFP (MOI=5) for four days. Cells were stained for UL44 (labeling the viral replication compartments) and analyzed by fluorescence microscopy. (e). THP1 macrophages, monocytes and monocytes overexpressing PDGFRα were infected with HCMV-GFP. At 7 dpi, viral supernatant was used to infect recipient wild-type fibroblasts. Forty-eight hours later, the percentage of GFP-positive recipient cells was determined by flow cytometry and used to calculate the number of plaque-forming units (PFU). n = 3 biological replicates. (f). Quantification of the number of viral particles within the cytoplasm of infected THP1 and THP1-PDGFRα monocytes or macrophages using HCMV-UL32-GFP (MOI=5) at 1.h.pi. (n Mac-PDGFRα=89, n Mac = 103, n mono =83, n mono-PDGFRα = 84). Viral particles were counted using FIJI image processing and statistics was performed using Poisson regression. (g). Light microscopy of THP1 and THP1-PDGFRα or THP1-THBD monocytes. (h). Flow cytometry analysis of cell surface staining of differentiation markers on THP-1 and THP-1–PDGFRα monocytes (upper panel) and THP1 macrophages (lower panel). Gating strategy is shown in Fig. S1a. For MHCII staining, monocytes treated with IFNγ are shown as positive control. (i). Differential gene expression analysis of Mass spectrometry data from THP1 and THP1–PDGFRα monocytes. Blue dots represent genes related to innate immunity (ISGs, compiled based on ^52^) and purple dots represent genes related to monocytes to macrophage differentiation (GO:0030225). Labeled genes are significantly differentially expressed (Padj < 0.05, fold change > 1) or mCherry and PDGFRA. n=3. (j). Differential gene expression analysis of RNA-seq data from THP1 and THP1–PDGFRα monocytes. Blue dots represent genes related to innate immunity (Interferon Stimulated Genes, ISGs, compiled based on^52^) and purple dots represent genes related to monocytes to macrophage differentiation (GO:0030225). (k) Flow cytometry analysis of PDGFRα surface expression in THP1 monocytes with induced expression of PDGFRα, compared to IgG control at 1 and 5 days post doxycycline treatment. Gating strategy is shown in Fig. S1a. (l). Flow cytometry analysis of THP1 monocytes and THP1 monocytes with induced expression of control or PDGFRα, infected with HCMV–GFP (MOI=5) at 3 dpi. The gate marks the productive, GFP-bright cell population. Gating strategy is shown in Fig. S1a. (m). Relative viral DNA in the supernatant of infected THP1 monocytes induced or constitutively expressed PDGFRα. Supernatant was collected at 8dpi. (n). Flow cytometry analysis of THP1-PDGFRα treated with DMSO or different concentrations of imatinib for 1h before infection with HCMV-GFP (MOI=5). Gating strategy is shown in Fig. S1a. (o). Flow cytometry analysis of NRP2 (left) or THBD (right) surface expression in THP-1 monocytes with induced expression of NRP2 or THBD, respectively, compared to the parental cells. Gating strategy is shown in Fig. S1a. (p). THBD and control cells were infected with HCMV-GFP. At 8 dpi, viral supernatant was used to infect recipient wild-type fibroblasts. Forty-eight hours later, the percentage of GFP-positive recipient cells was determined by flow cytometry and used to calculate the number of plaque-forming units (PFU). n = 2. (q). Flow cytometry analysis of THP1 overexpressing THBD infected with HCMV–GFP (MOI=5) at 3 dpi showing HCMV-GFP level versus THBD surface level. Gating strategy is shown in Fig. S1a. (r). Differential gene expression analysis of RNA-seq data from THP1 mono overexpressing mCherry control or THBD. Blue dots represent genes related to innate immunity (Interferon Stimulated Genes, ISGs, compiled based on ^52^) and purple dots represent genes related to monocytes to macrophage differentiation (GO:0030225). (s). Relative viral DNA on infected primary monocytes overexpressing THBD or PDGFRα at 3 and 11 dpi. n=2.

**Fig. S5.**
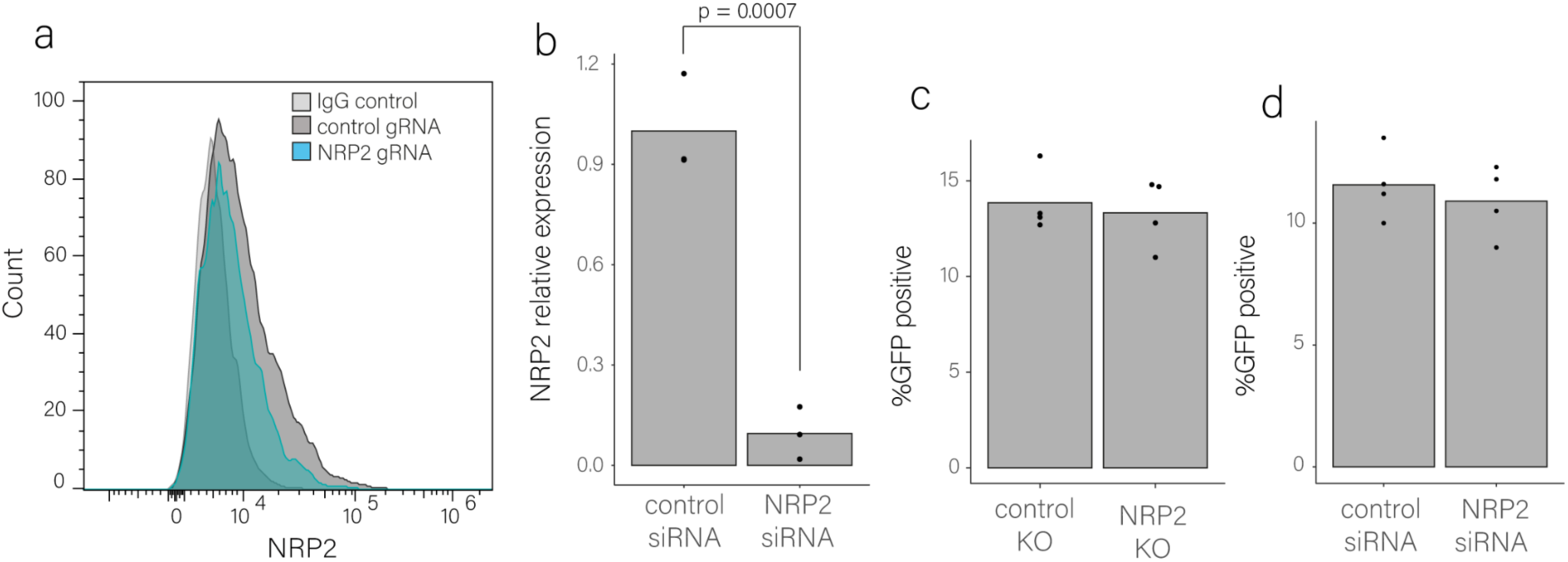
(a). Cell surface staining with NRP2 or IgG control of THP1 macrophages treated with CRISPR knockout against NRP2 or control. Cells were analyzed by Flow cytometry. Gating strategy is shown in Fig. S1a. (b). Relative expression level of NRP2, measured by real-time PCR in THP1 macrophages treated with NRP2 siRNA compared to control siRNA at the time of HCMV infection. p-value was calculated using a two-sided student t-test. n=3. (c). Quantification of flow cytometry analysis of THP1 macrophages with NRP2 or control CRISPR knockout, infected with HCMV-GFP (MOI=5). Analysis was performed at 3 dpi, p-value was calculated using a two-sided student t-test. n=4. (d). Quantification of flow cytometry analysis of THP1 macrophages, transfected with NRP2 and control siRNA two days before infection with HCMV-GFP (MOI=5). Analysis was performed at 3 dpi, p-value was calculated using a two-sided student t-test. n=4.

**Fig. S6.**
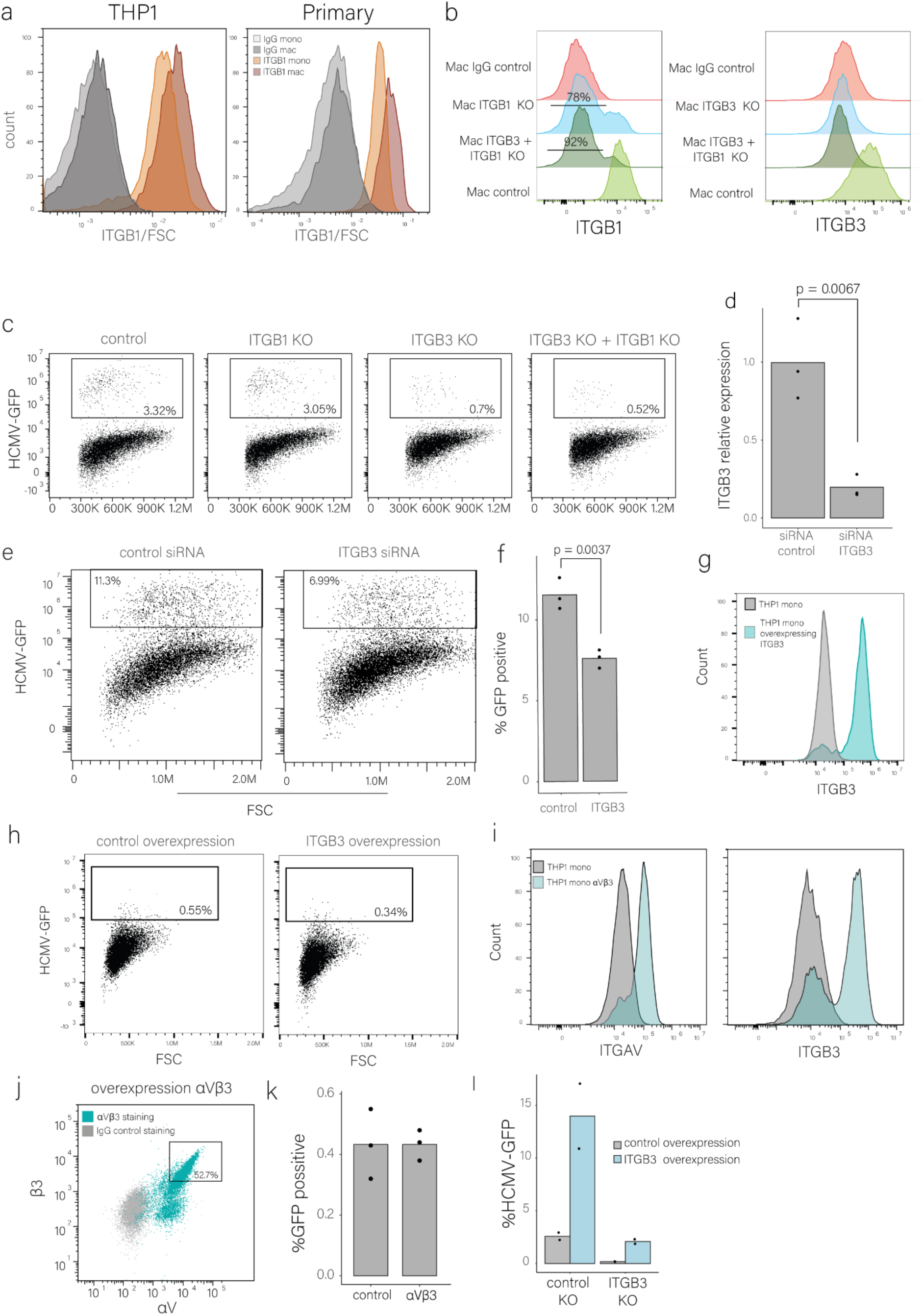
(a). Cell surface levels of ITGB1 in primary and THP1 monocytes and macrophages that are presented in Fig. 6c, normalized to FSC (as a measure for the size of the cells). (b). Flow cytometry analysis of cell surface staining of THP1 macrophages with CRISPR knockout of β1 (left) and/or β3 (right) versus control knockout. Gating strategy is shown in Fig. S1a. (c). Flow cytometry analysis of control, ITGB3, ITGB1 and ITGB3 + ITGB1 knockout (KO) in THP1 macrophages infected with HCMV–GFP (MOI=5). Cells were analyzed at 3 dpi. Gating strategy is shown in Fig. S1a. (d). real-time PCR analysis of THP1 macrophages treated with siRNA against ITGB3 or control at 0 hpi. P-value was calculated using a two-sided student t-test. n=3. (e). Flow cytometry analysis of THP1 macrophages, transfected with ITGB3 or control siRNAs and infected with HCMV–GFP (MOI=5). Cells were analyzed at 3 dpi. Gating strategy is shown in Fig. S1a. (f). Quantification of the FACS replicates of Fig. S6e. P-value was calculated using a two-sided student t-test. n=3. (g). Flow cytometry analysis of ITGB3 surface expression in THP-1 monocytes with induced expression of ITGB3 compared the parental cells. Gating strategy is shown in Fig. S1a. (h). Flow cytometry analysis of infected THP1 monocytes overexpressing ITGB3, compared to mCherry control. Overexpression was induced for 24 hours using doxycycline prior to HCMV-GFP (MOI=5) infection. Cells were analyzed at 3 dpi. Gating strategy is shown in Fig. S1a. (i). Cell surface staining of THP1 monocytes overexpressing (OE) αVβ3 compared to parental cells. Cells were stained for either αV or β3 in the αVβ3 overexpressing cells. Gating strategy is shown in Fig. S1a. (j). Cell surface staining of THP1 overexpressing (OE) αVβ3 compared to IgG control. Cells were stained for both αV and β3 in the αVβ3 overexpressing cells. The gate marks the double positive cells which were sorted prior to HCMV infection. Gating strategy is shown in Fig. S1a. (k). Quantification of flow cytometry analysis of infected THP1 monocytes overexpressing ITGB3 and ITGAV (αvβ3), compared to mCherry control presented in figure 6g. Overexpression was induced for 24 hours using doxycycline prior to HCMV-GFP (MOI=5) infection. Cells were analyzed at 3 dpi. n=3. (l). Quantification of flow cytometry results of infected THP1 macrophages with HCMV-GFP (MOI=5) overexpressing ITGB3 or mCherry control in the background of ITGB3 or control knockout. Cells were analyzed at 3 dpi. Gating strategy is shown in Fig. S1a.

**Fig. S7.**
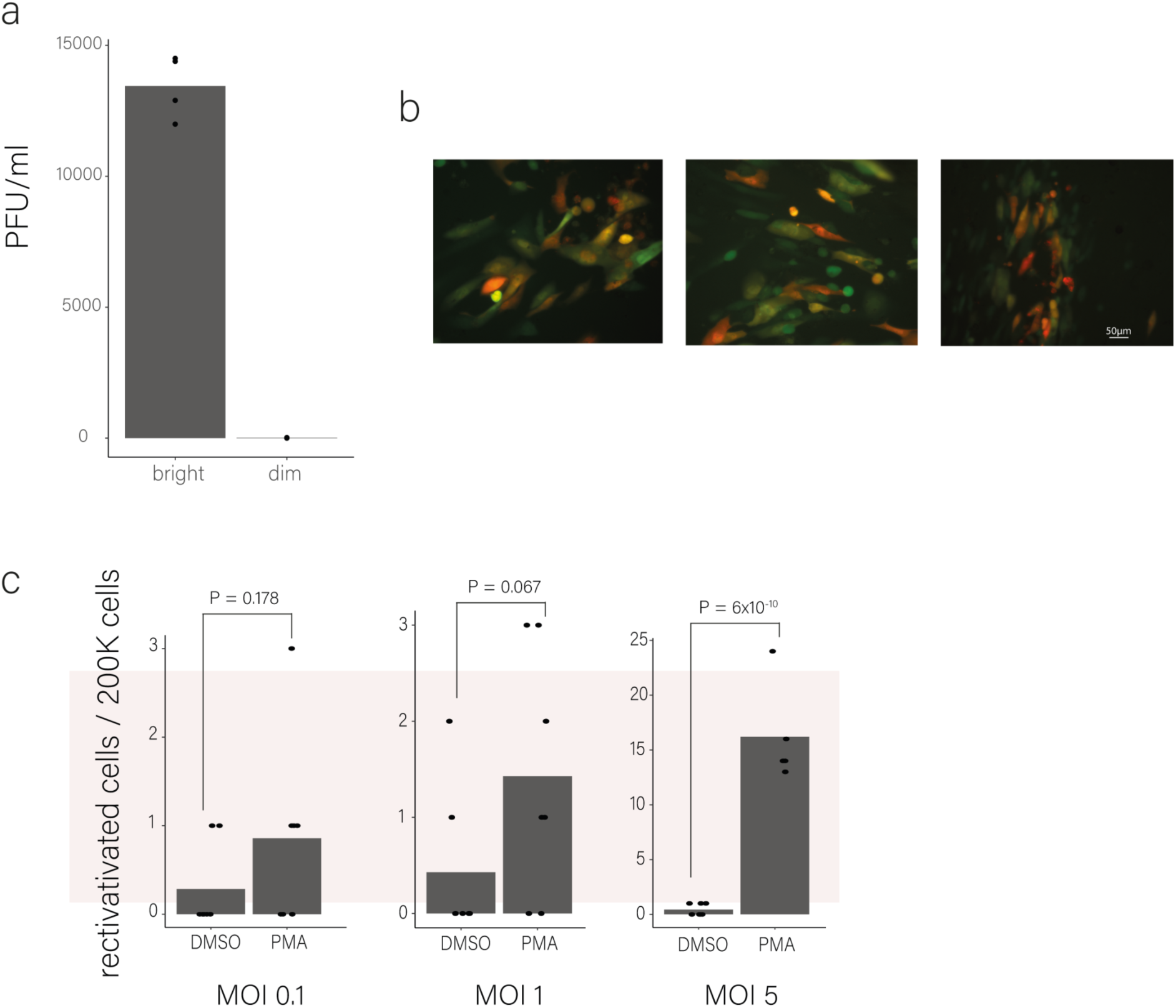
(a). Measurement of infectious virus in supernatant from FACS-sorted IE-bright and IE-dim infected inducible PDGFRα THP1 monocytes, collected at 8dpi, PFU, plaque-forming units. n = 4. (b). Images of infectious centers from reactivated inducible PDGFRα THP1 monocytes, showing IE gene expression (Green), and late gene expression (Red). (c). Reactivation levels of HCMV from the sorted dim populations presented in (7f), cells were treated with PMA or DMSO as a control at 7 d.p.i and GFP-positive cells were counted three days later. Statistics performed using Poisson regression.

